# Microglia regulate GABAergic neurogenesis in prenatal human brain through IGF1

**DOI:** 10.1101/2024.10.19.619180

**Authors:** Diankun Yu, Samhita Jain, Andi Wangzhou, Stacy De Florencio, Beika Zhu, Jae Yeon Kim, Jennifer Ja-Yoon Choi, Mercedes F Paredes, Tomasz J Nowakowski, Eric J Huang, Xianhua Piao

## Abstract

GABAergic neurons are an essential cellular component of neural circuits. Their abundance and diversity have enlarged significantly in the human brain, contributing to the expanded cognitive capacity of humans. However, the developmental mechanism of the extended production of GABAergic neurons in the human brain remains elusive. Here, we use single-cell transcriptomics, bioinformatics, and histological analyses to uncover microglial regulation of the sustained proliferation of GABAergic progenitors and neuroblasts in the human medial ganglionic eminence (hMGE). We show that insulin-like growth factor 1 (IGF1) and its receptor IGR1R as the top ligand-receptor pair underlying microglia-progenitor communication in the prenatal human brain. Using our newly developed neuroimmune hMGE organoids, which mimics hMGE cytoarchitecture and developmental trajectory, we demonstrate that microglia-derived IGF1 promotes progenitor proliferation and the production of GABAergic neurons. Conversely, IGF1-neutralizing antibodies and *IGF1* knockout human embryonic stem cells (hESC)-induced microglia (iMG) completely abolished iMG-mediated progenitor proliferation. Together, these findings reveal a previously unappreciated role of microglia-derived IGF1 in promoting proliferation of neural progenitors and the development of GABAergic neurons.

## Main Text

In the adult human neocortex, about 25-50 percent of cortical neurons are γ-aminobutyric acid-containing (GABAergic) inhibitory interneurons ^1–3^. They serve as the principal source of cortical inhibitory input and provide a crucial role in maintaining the excitation-inhibition balance and functional rhythms in the brain ^4,5^. Disruptions in interneuron numbers and functions have been implicated in various neurological and psychiatric disorders including autism spectrum disorders (ASD) ^6,7^, epilepsy ^8^, and schizophrenia ^9^. In the prenatal human brain, GABAergic interneurons are generated in the ganglionic eminences (GEs) of the ventral telencephalon ^10,11^. The medial ganglionic eminence (MGE) gives rise to most of the parvalbumin-positive (PV^+^) and somatostatin-positive (SST^+^) cortical interneurons ^6^. The human MGE (hMGE) possesses several unique features, distinct from other species, to meet the needs of a drastically expanded cerebral cortex. First, neurogenesis in the hMGE is most active during the second and third trimesters ^12^. This active neurogenesis is followed by extensive migration of GABAergic neurons around the periventricular zone in the neonatal stage until at least 6 months postnatally ^13^. Second, young GABAergic neuroblasts are organized as DCX^+^ cells-Enriched Nests (DENs) that have sustained proliferation up to the early postnatal stage. GABAergic neuroblasts within DENs are surrounded by NESTIN^+^ and SOX2^+^ radial glia that extend from the ventricular zone (VZ) and inner subventricular zone (iSVZ) to the outer SVZ (oSVZ) ^10,12^. Third, the DCX^+^SOX2^+^ neuroblasts inside DENs exhibit regional differences in their proliferation potentials, with those at the edge of DENs showing a higher Ki-67 labeling index ^12^. While these observations support that DCX^+^ neuroblasts in DENs have the capacity to undergo sustained neurogenesis in the hMGE ^10,12^, it remains largely unclear what are the external environmental cues that regulate hMGE neurogenesis.

Microglia, the specialized tissue-resident macrophages in the central nervous system, are the primary immune cells in the brain parenchyma. Originated from the yolk sac and entering the brain during early gestation ^14^, microglia have been shown to be involved in brain development, including neurogenesis ^15^, angiogenesis ^16,17^, neuronal survival ^18^, myelination ^19–22^, synaptogenesis ^23^, and synaptic pruning ^24–28^. It has been shown that microglia regulate PV^+^ interneuron development and positioning ^29^, and lead to PV^+^ interneuron deficit in setting of maternal immune activation in the mouse brain ^30^. However, it remains unclear whether and how microglia could impact the development of GABAergic neurons in the human brain. Here, we show that during the late second trimester, the iSVZ and oSVZ of hMGE are populated with microglia that are near radial glia and Ki-67^+^ proliferating neuroblasts at the periphery of DENs. Single-nucleus transcriptomics studies of human brains from the late second trimester to the early postnatal stage uncover insulin-like growth factor 1 (IGF1) and its receptor IGF1R as the top candidate pathway in mediating the communications between microglia and interneuron progenitors. Indeed, further studies, using human MGE neuroimmune organoids (MGEO) where hESC-derived microglia (iMG) were transplanted into human pluripotent cells (hPSC)-derived MGE organoids, support that microglia-derived IGF1 promotes progenitor proliferation and interneuron production.

### Microglia intimately associate with radial glia and neuroblasts in the hMGE

To study microglial function in hMGE development, we first surveyed their temporal and spatial distributions in the hMGE of neurotypical human postmortem brains from gestational weeks (GW) 15 to GW41 by IBA1 immunohistochemistry (IHC) (Fig. 1a-d). At GW15-17, microglia were sparsely sprinkled throughout the hMGE (Fig. 1a, d). By GW22-24, however, there was a clear locational preference, with most microglia seen in the iSVZ and oSVZ (Fig. 1b, d). Interestingly, microglia in the oSVZ accumulated in the area outside of DENs (oDENs) and encased the outer rim of DENs with rare microglial processes extending into the inside of DENs (iDENs) (Fig. 1b, d). By GW39-41, the density of microglia in the iSVZ and oSVZ of hMGE remained high, and more microglial processes could be identified in the iDENs compared to those in the late second trimester (Fig. 1c, d).

**Fig. 1.**
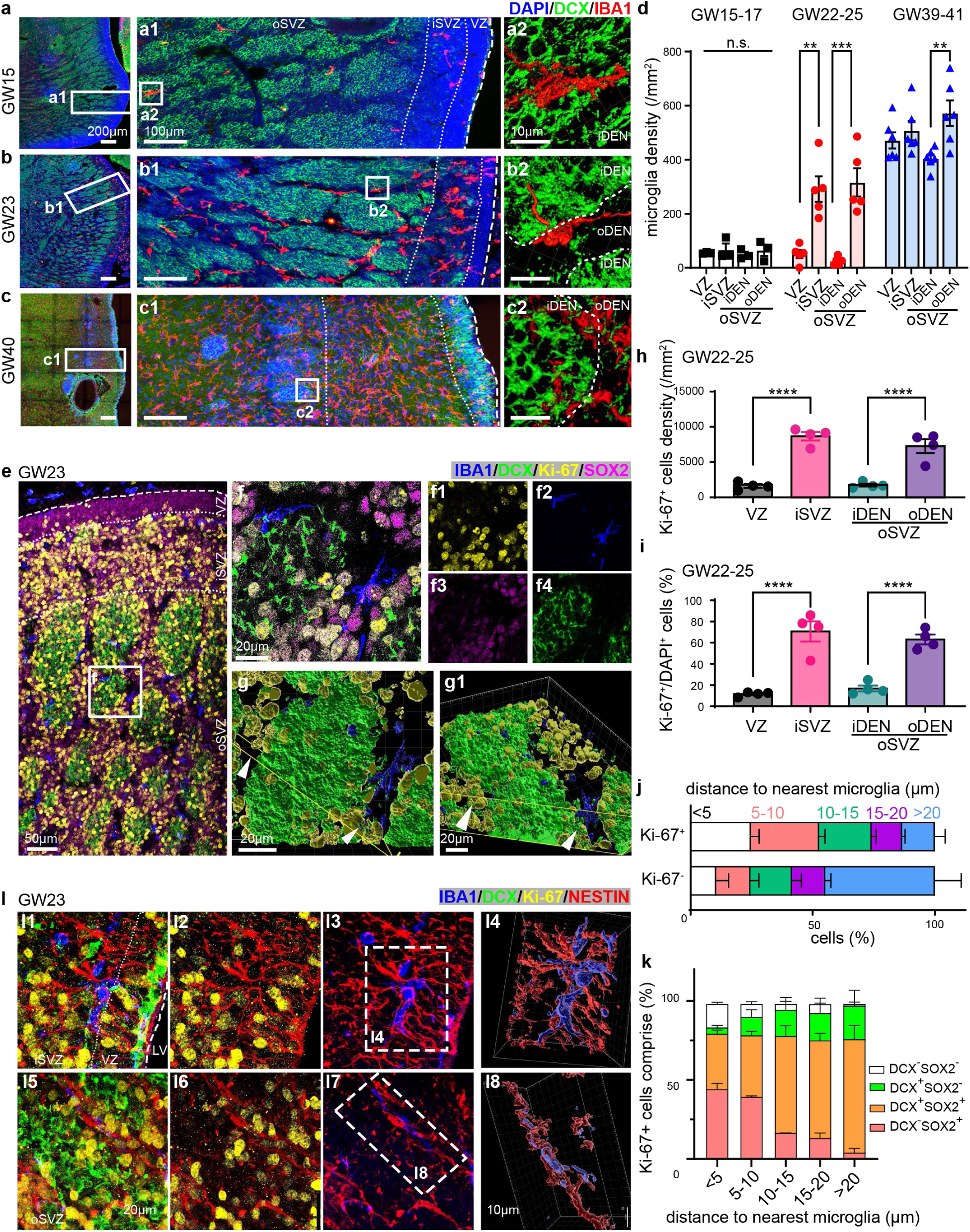
Microglia are in proximity with MGE progenitors in the developing human brains. **a-d,** IHC images and bar graphs showing microglia in the developing hMGE at GW15, GW23 and GW40. Microglia are highly concentrated in the iSVZ and the oDENs of the oSVZ at GW22-GW24. **e-f**, IHC images showing the spatial relationship between Ki-67^+^ proliferating progenitors and microglial in GW23 hMGE sections. **g**, Three-dimensional reconstruction revealing the close proximity between microglia and Ki-67^+^ proliferating progenitors in the oSVZ of GW23 hMGE. Arrowheads point to the Ki-67^+^ clusters near microglia in the oSVZ. **h**-**i**, Bar graph showing the density and percentage of Ki-67^+^ progenitors in GW22-GW24 hMGE. **j**, The percentage of Ki-67^+^ and Ki-67^−^ cells as their distance to microglia increases in the oSVZ of hMGE. **k**, Cell composition analysis of Ki-67^+^ progenitors showing Ki-67^+^ cells in proximity to microglia comprise of SOX2^+^DCX^−^ radial glia and SOX2^+^DCX^+^ neuroblasts. **l**, IHC images and three-dimensional reconstruction indicating that microglia closely contact NESTIN^+^ projections in GW23 hMGE. For statistics, N=3, 5 and 6 in (d); N=4 in (h) and (i); N=4 for in (j) and (k). ** indicates P<0.01, **** indicates P<0.001, n.s. indicates non-significant; some non-significant comparisons are not labelled; one-way ANOVA test and post-hoc Bonferroni’s test for (d), (h) and (i); two-way repeated measures ANOVA for (j) and (k) showed significant interaction effect; Data in (d), (h), (i), (j) and (k) were shown as means ± standard error of the mean (SEM).

To further examine the spatial relationship between microglia and proliferating cells in the hMGE during GW22-24, we performed a combined IHC with IBA1 and Ki-67, a marker for cell proliferations. We saw that microglia mostly resided in regions of more abundant proliferative progenitors (Fig. 1e-i). Specifically, many microglia were embedded within Ki-67^+^ proliferating progenitors in the iSVZ and oDENs of the oSVZ (Fig. 1e, f). Through three-dimensional (3D) reconstruction of DENs with IMARIS software, we found that Ki-67^+^ cells were largely clustered around microglia on the outside and the border of DENs, with a gradual decreased ratio of Ki-67^+^ cells over Ki-67^−^ cells as the distance to microglia increased (Fig. 1g, j). Furthermore, Ki-67^+^ progenitors in proximity to microglia were largely SOX2^+^ progenitors, including SOX2^+^DCX^−^ radial glia and SOX2^+^DCX^+^ neuroblasts (Fig. 1e-g, k). To further document the spatial relationship between microglia and MGE progenitors, we performed IHC of IBA1, DCX, NESTIN, and Ki-67. We found that microglia were in close contact with and often followed along NESTIN^+^ radial glial fibers (Fig. 1l). In the iSVZ, microglia were mostly ramified and intermingled with multiple NESTIN^+^ fibers (Fig. 1l1-l4). Interestingly, most microglia in the oSVZ exhibited polarized morphology along the NESTIN^+^ fibers, with their processes largely aligning and closely contacting with those fibers (Fig. 1l5-l8).

Distinct from what was observed in the hMGE, the density of microglia was significantly higher in the VZ of mouse MGE at embryonic day (E) 14.5 (Extended Data Fig. 1a-c) and 16.5 (Extended Data Fig. 1d-f), coincident with higher density of active proliferating progenitors in the VZ of mouse MGE (Extended Data Fig. 1g-h). This contrast between mouse and human MGE could highlight a potential species-specific mechanism of interneuron development.

### Single-cell transcriptomics reveal microglia-mediated mechanisms that promote interneuron development

To identify molecular underpinnings of microglial regulation of extended neurogenesis during human interneuron development, we generated a single-nucleus RNA-sequencing (snRNA-seq) dataset of late embryonic stage (GW22-GW30) to perinatal stage (postnatal week (PW) 2-PW 3) using droplet-based 10X Genomics platform (N=6 donors, Fig. 2a and Extended Data Table 1). We focused on human tissues that contained the ganglionic eminences, adjacent periventricular regions (containing the Arc, an area of the subventricular zone composed of migratory interneurons^13^), and cortical regions where mature interneurons reside (Fig. 2a and Extended Data Table 1). Neurons are overrepresented in snRNA-seq ^31^. To better capture both neurons and glia, we adapted flow cytometry-based strategies to enrich microglia through PU.1^+^, and oligodendrocyte lineage cells and MGE progenitors through OLIG2^+^, in addition to DAPI^+^ nuclei ^32^ (Fig. 2b and Extended Data Table 1).z After several quality control (QC) steps and doublets removal (details in Methods), we recovered 124,411 nuclei with a median of 7,341 unique molecular identifiers and 2,925 genes per cell. We then performed principal component analysis of normalized reads counts followed by uniform manifold approximation and projection (UMAP) using Seurat-v5 ^33^. We identified eleven main cell clusters from our transcriptomic data profiles based on the expression of canonical maker genes (Fig. 2c and Extended Data Fig. 2 and 3). Notably, our results contained 17,342 cells in the microglia cluster and 17,667 cells in the cortical GABAergic interneuron cluster that includes GE progenitors, young GABAergic interneurons, and mature GABAergic interneurons (Extended Data Fig. 2).

**Fig. 2.**
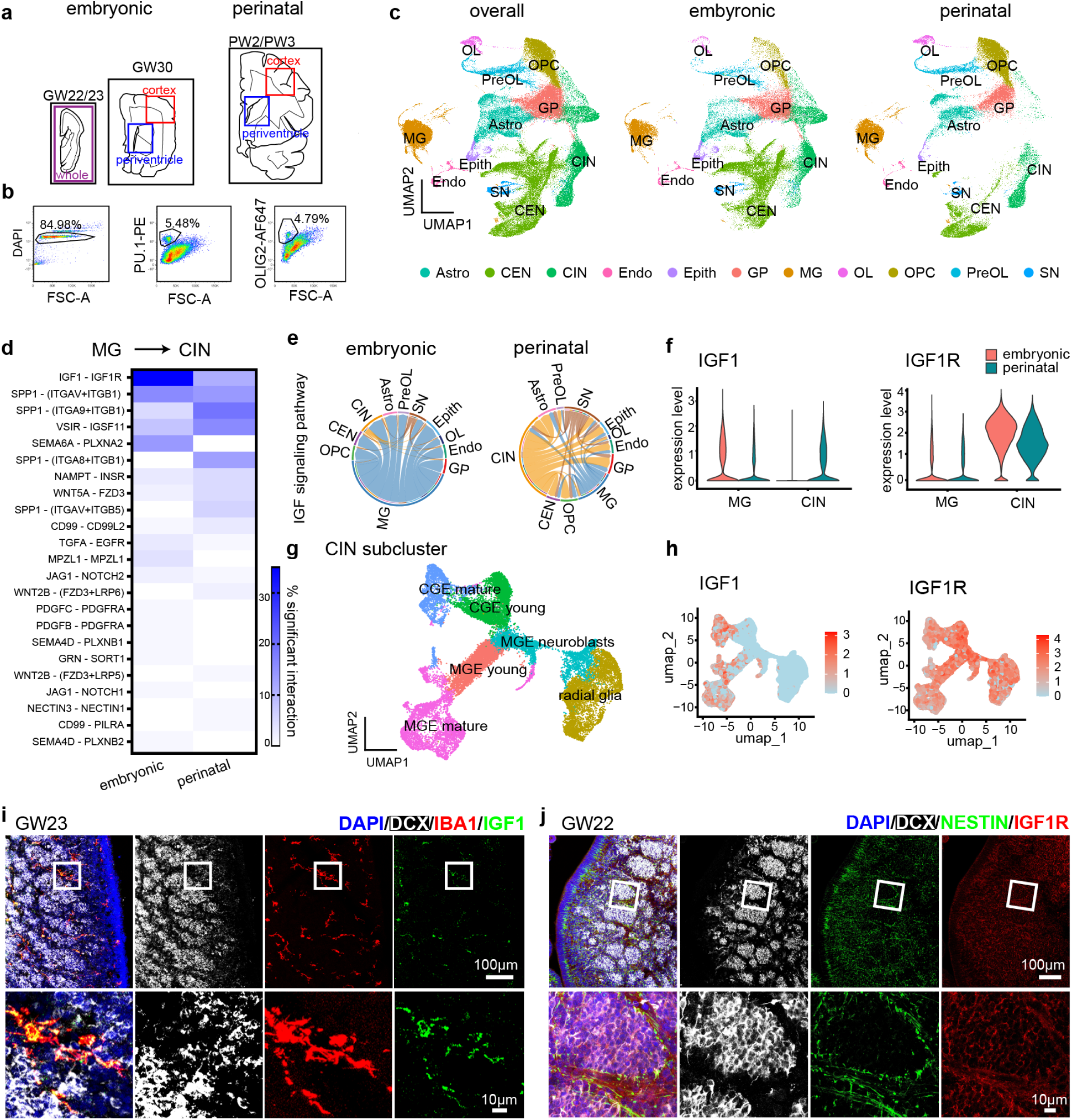
Transcriptomic profiling and cell-cell interaction analysis uncover IGF1-IGF1R as the top potential signaling pathway mediating the communication between interneuron progenitors and microglia in the developing human MGE. **a,** Diagrams showing the brain regions used for snRNA-seq experiment. **b**, Flow cytometry strategies for unenriched DAPI^+^ nuclei and the enrichment of PU.1^+^ nuclei and OLIG2^+^ nuclei. **c**, UMAP plots showing the eleven cell types identified. Astro: astrocytes; CEN: cortical excitatory neurons; CIN: cortical GABAergic interneurons; Endo: endothelial cells; Epith: epithelial cells; GP: glia progenitors; MG: microglia; OL: oligodendrocytes; OPC: oligodendrocyte precursor cells; PreOL: pre-myelinating oligodendrocytes; SN: subpallial neurons. **d**, Heatmap plot showing the significant ligand-receptor pairs that potentially mediate microglial regulation of interneuron development at the different developmental stages. **e**, Chord plots of IGF signaling pathways at the embryonic and perinatal stages. **f**, Violin plots of IGF1 and IGF1R expression by microglia and interneurons at the embryonic and perinatal stages. **g**, UMAP plots of subtypes of cortical GABAergic interneurons, including radial glia, MGE neuroblasts, MGE-derived young interneurons (MGE young), MGE-derived mature interneurons (MGE mature), CGE-derived young interneurons (CGE young), and CGE-derived mature interneurons (CGE mature). **h**, Feature plots showing the expression pattern of IGF1 and IGF1R in interneuron subtypes. **i**, IHC showing the specific expression of IGF1 in microglia in the developing hMGE. **j**, IHC showing the expression pattern of IGF1R in the developing hMGE.

To predict cell type-specific interactions between microglia and other cell types in the developing hMGE, we leveraged a highly curated database of receptor-ligand interactions to predict possible interactions among different types of cells (Extended Data Fig. 4) ^34^. This approach also revealed development-related pathways by comparison analysis of cell-cell communications at the late embryonic and perinatal stages (Extended Data Fig. 5). To gain insight into the interaction involved in microglial regulation of cortical interneurons, we extracted those ligand-receptor pairs with statistical significance (Fig. 2d). Among them, IGF1 and IGF1R was the ligand-receptor pair with the highest communication probability and predominantly presented in the embryonic stages (Fig. 2d). Chord plots, violin plots and feature plots showed that IGF1 was predominantly derived from microglia during the embryonic stage, while it was also generated by interneurons at the perinatal stage (Fig. 2e, f, and Extended Data Fig. 6). To clarify which interneuron population expresses IGF1, we conducted interneuron subcluster analysis and identified unique populations, namely radial glia, MGE neuroblasts, MGE-derived young interneurons, MGE-derived mature interneurons, caudal GE (CGE)-derived young interneurons, and CGE-derived mature interneurons, according to unbiased clustering and canonical markers (Fig. 2g and Extended Data Fig. 7). We observed that IGF1 was mainly expressed by mature interneurons, but barely detected in interneuron progenitors, including radial glia and MGE neuroblasts (Fig. 2h). These results indicated that microglia were the key source of IGF1 during interneuron proliferation in the developing hMGE. Accordingly, IGF1R is highly expressed in interneurons, particularly at the embryonic stage (Fig. 2e, f, and Extended Data Fig. 6). Subcluster analysis further confirmed the high expression level of IGF1R in interneuron progenitors and young interneurons (Fig. 2g, h). Indeed, our IHC showed that, in GW22-24 hMGE, IGF1 was specifically expressed in microglia (Fig. 2i), whereas IGF1R was widely expressed in progenitors in the hMGE (Fig. 2j). Taken together, our data support that IGF1 is primarily derived from microglia during the embryonic state and has a potential to support interneuron development in the hMGE.

### hPSC-derived MGE organoids (MGEOs) recapitulate hMGE development

To study the role of human microglia in hMGE development, we established microglia-containing MGEO by transplanting hESC-induced iMG to hPSC (including both human induced pluripotent stem cells (hiPSC) and hESC)-derived MGE-oriented organoids ^35,36^ (Fig. 3a). Our MGEOs had robust NKX2.1^+^ and Ki-67^+^ VZ-like rosettes at 6 weeks old (Fig. 3b) and demonstrated sequential expression of markers for MGE progenitors and interneurons, including DLX2, SOX2, LHX6, DCX, GAD67, NeuN, SST, and PV (Fig. 3c-f, and Extended Data Fig. 8a), confirming the MGE identity and generation of GABAergic interneurons. Notably, Ki-67-expressing cells are distributed in the center of VZ-like rosettes as well as the edge and neighboring SVZ-like regions of rosettes at 6 weeks old (Fig. 3b and Extended Data Fig. 8b). IHC analysis showed that Ki-67^+^ progenitors were comprised of both SOX2^+^DCX^−^ radial glia-like cells and SOX2^+^DCX^+^ neuroblasts in 6-week-old MGEOs (Extended Data Fig. 8b), recapitulating spatial organization radial glia and proliferating neuroblasts in the developing hMGE. Interestingly, we observed clusters of DCX^+^ neuroblasts sprinkled with Ki-67^+^ cells in 24-week-old MGEOs, which could represent DEN-like clusters (Fig. 3g). Taken together, our data supports that MGEOs can serve as a model to study hMGE development.

**Fig. 3.**
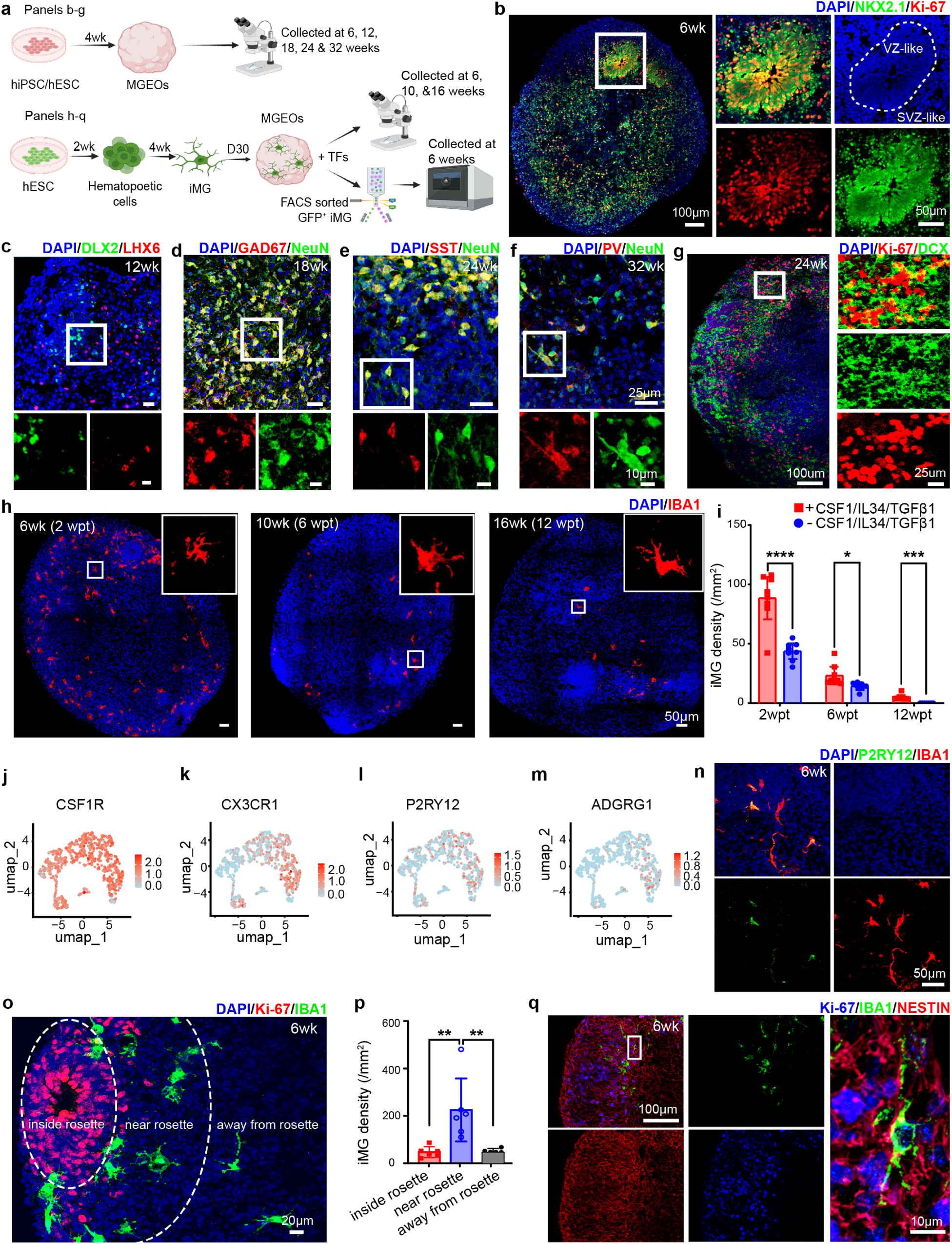
MGEO models hMGE development and interaction between microglia and hMGE progenitors. **a**, Schematic diagram of experimental flow. TFs includes CSF1, IL34, and TGFβ1. **b**, MGEO containing “rosette”-like proliferating centers, recapitulating the VZ- and SVZ-like areas of the hMGE. **c**-**f**, Sequential expression of markers for MGE progenitors and MGE-derived interneurons that mimic the temporal progression of hMGE development. **g**, Clusters of DCX^+^ neuroblasts sprinkled with Ki-67^+^ cells noted in 24-week-old organoids, resembling DEN-like structures in the hMGE. **h**-**i**, iMG survive up to 12 weeks post iMG transplantation (wpt) in MGEO. **j**-**m**, Feature plots of microglial homeostatic markers expressed in FACS-isolated iMG from 6-week-old MGEOs. **n**, IHC images showing the expression of P2RY12 in iMG. **o**-**p**, IHC images and quantification showing the distribution of iMG around rosettes. **q**, IHC images showing microglia closely interact with NESTIN^+^ projections in organoids. For statistics, N=8,8,8,7,8,7 in each group in (i) and N= 6 in (p). * indicates P<0.05, ** indicates P<0.01, ***P<0.001, ****P<0.0001, unpaired t-test in (i); one-way ANOVA and post-hoc Bonferroni’s test in (p). Data in (i) and (p) were shown as means ± SEM.

We transplanted iMG into MGEOs at 4-week-old, mimicking the stage when microglia are detected in developing human brains at GW4.5 ^14,37^ (Fig. 3a). The addition of recombinant human colony stimulating factor 1 (CSF1), interleukin 34 (IL34), and transforming growth factor beta 1 (TGFβ1) to the culture medium significantly improved iMG survival up to 12 weeks post transplantation (wpt) (Fig. 3h, i). The transplanted iMG demonstrated similar morphologies and transcriptomics with human primary microglia (Fig. 3h, j-n). Importantly, they expressed homeostatic microglia markers such as CSF1R (Fig. 3j), CX3CR1 (Fig. 3k), P2RY12 (Fig. 3l, n), and particularly ADGRG1 (Fig. 3m), one of the few genes that define yolk sac-derived “true” microglia ^38^ (Fig. 3m). As was seen in GW22-GW24 hMGE, we observed iMG mostly avoided the VZ-like rosettes but accumulated in the SVZ-like neighboring regions in the MGEOs (Fig. 3o, p), and intimately interacted with NESTIN^+^ projections (Fig. 3q).

### Microglia promote MGE proliferation and interneuron production

To investigate microglial regulation of interneuron development, we conducted single-cell RNA-seq of 6-week-old MGEOs with and without transplanted iMG at 4-week-old (Fig. 4a). Based on unbiased clustering and canonical marker genes, we identified clusters of radial glia, neuroblasts, young MGE-derived GABAergic interneurons, and a small group of CGE-like cells (Fig. 4b and Extended Data Fig. 9). We discovered a significant increase in the proportion of radial glia as well as a significant higher percentage of Ki-67^+^ cells in MGEOs transplanted with iMG (Fig. 4c, d), indicating a role of iMG in promoting progenitor proliferation. Differentially expressed genes (DEGs) analysis showed that upregulated genes in MGEOs transplanted with iMG were enriched in pathways and gene ontologies (GOs) related to cell mitosis (Fig. 4e, f, and Extended Data Fig. 10). The downregulated genes in MGEOs with iMG were largely enriched in pathways and GOs related to oxidative stress (Fig. 4e, f, and Extended Data Fig. 10), as is shown in previous studies ^39,40^. Additionally, upregulated genes in radial glia were also significantly enriched in Gene Expression Omnibus (GEO) terms related to IGF1R downstream signaling (Fig. 4e, f), such as CCND1, TMPO, RRM2, and MCM3, in MGEOs with iMG, suggesting a role of IGF1-IG1R in microglial regulation of interneuron progenitor proliferation.

**Fig. 4.**
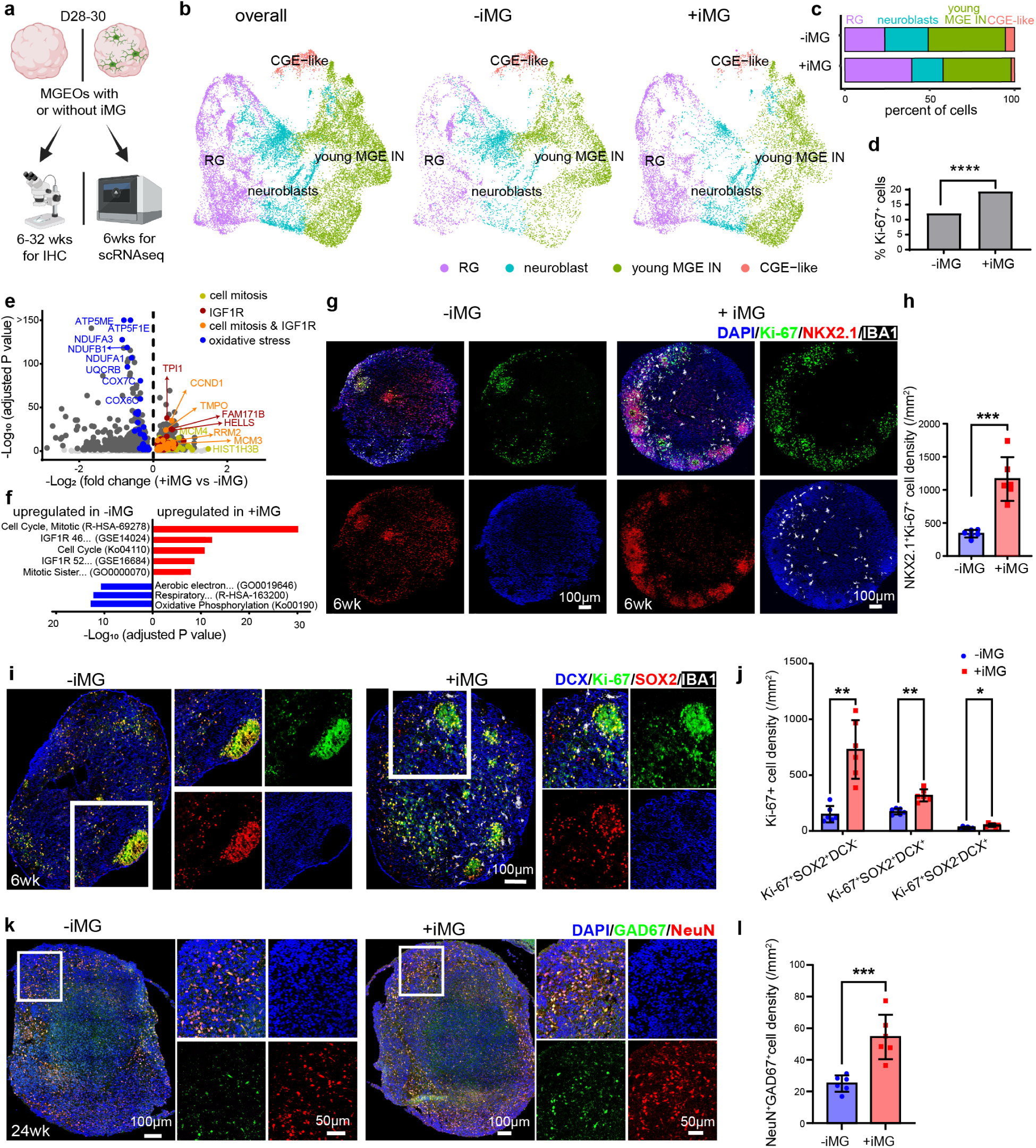
Microglia promote MGE proliferation and interneuron production. **a**, Experimental diagrams. **b**, UMAP plots showing cell type clusters from 6-week-old MGEOs with and without iMG. RG: radial glia; young MGE IN: young MGE-derived GABAergic interneurons. **c**, Bar graph showing the proportion of RG is significantly increased in MGEOs transplanted with iMG (P<0.0001). **d**, Bar graph showing the percentage of Ki-67^+^ nuclei is significantly increased in MGEOs with iMG. **e**, Volcano plot showing DEGs in RG of MGEO with and without iMG. **f**, Upregulated DEGs in MGEO with iMG are enriched in pathways and GOs related to cell mitosis and IGF1R signaling pathway; upregulated DEGs in MGEO without iMG are enriched in pathways and GOs related to oxidative stress. IGF1R 46: IGF1R drug inhibition 46 (Kinase Perturbations from GEO down, GSE14024); IGF1R 52: IGF1R knockdown 52 (Kinase Perturbations from GEO down, GSE16684); Mitotic Sister: Mitotic Sister Chromatid Segregation (GO:0000070); Aerobic electron: Aerobic Electron Transport Chain; Respiratory: Respiratory Electron Transport, ATP Synthesis By Chemiosmotic Coupling, Heat Production By Uncoupling Proteins (R-HSA-163200); **g**-**h**, IHC images and bar graph showing that the presence of iMG increases the density of NKX2.1^+^Ki-67^+^ progenitors in 6-week-old MGEO. **i**-**j**, IHC images and bar graph showing the presence of iMG leads to an increased density of SOX2^+^DCX^−^ radial glia in 6-week-old MGEOs. **k**-**l**, IHC images and bar graph showing that the presence of iMG increases the density of NeuN^+^GAD67^+^ interneurons in 18-24-week-old MGEOs. For statistics, * indicates P<0.05; ** indicates P<0.01, *** indicates P<0.001, **** indicates P<0.0001; N=6 in (h); N=6 in (j); N=6 in (l); χ^2^ test, 2221 (RG cell number)/9456(overall cell number) vs 4605/11680, χ^2^=607.1, P<0.0001 in (c); χ^2^ test, 1146(Ki-67^+^ cell number)/9456(overall cell number) vs 2269/11680, χ^2^=206.0 in (d); unpaired t-test in (h) and (l). Two-way ANOVA and post-hoc Bonferroni’s test in (j); Data in h, j and l were shown as means ± SEM.

We next conducted IHC to characterize the effect of iMG on interneuron development in MGEOs. Using Ki-67 and NKX2.1 as markers, we observed that the density of proliferating MGE progenitors was significantly increased in 6-week-old MGEOs transplanted with iMG (Fig. 4g, h, and Extended Data Fig. 11). Further analysis revealed that the presence of iMG significantly increased the density of proliferating radial glia (Ki-67^+^SOX2^+^DCX^−^) and neuroblasts (Ki-67^+^SOX2^+^DCX^+^) (Fig. 4i, j). Excitingly, we found that iMG transplantation significantly increased the density of NeuN^+^GAD67^+^ mature interneurons in 18-24-week-old organoids (Fig. 4k, l), as well as a trend of increased NeuN^+^PV^+^ and NeuN^+^SST^+^ MGE-derived subtypes of interneurons in 24- and 32-week-old organoids (Extended Data Fig. 12). Taken together, our data support that microglia promote the proliferation and generation of MGE-derived interneurons.

### Microglia derived IGF1 promotes MGE progenitor proliferation in the MGEOs

We next investigated whether microglia exert their function in hMGE development through IGF1 using our neuroimmune MGEOs (Fig. 5a). Importantly, we showed that IGF1 was specifically detected in iMG, while IGF1R was highly expressed by NESTIN^+^ radial glia in 6-week-old MGEOs (Fig. 5b, c), as observed in hMGE (Fig. 2i, j).

**Fig. 5.**
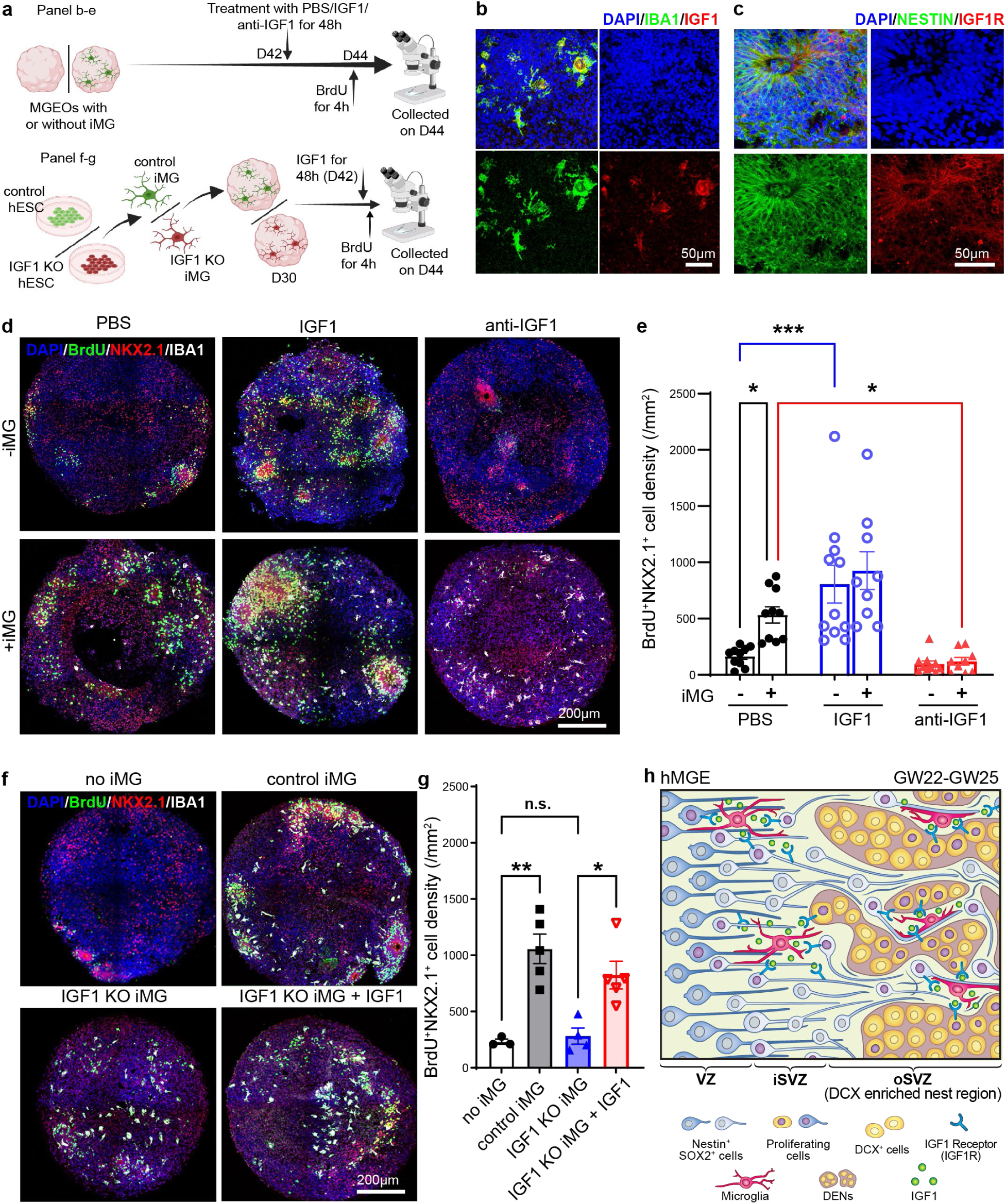
Microglia-derived IGF1 promotes MGE proliferation in MGEO. **a**, Experimental schematics. **b**, IHC images indicating the specific expression of IGF1 in iMG in 6-week-old MGEOs. **c**, IHC images showing the expression of IGF1R in progenitors in the rosettes of 6-week-old MGEOs. **d**-**e**, IHC images and bar graphs showing that IGF1 promotes MGE progenitor proliferation in MGEOs without iMG to the level seen in MGEOs with iMG, and that IGF1-neutralizing antibody treatment abolishes iMG-mediated progenitor proliferation. **f**-**g**, IHC images and bar graphs showing that the inability of IGF1 KO iMG to promote MGEO progenitor proliferation, which can be rescued by the addition of recombinant IGF1 treatment. **h**, Graphic abstract showing microglial regulation of interneuron progenitor proliferation via IGF1 in the hMGE. For statistics, N= 10, 11, 9 for MGEO -iMG, N=10, 9, 9 for MGEO +iMG in (e); N=3,5,5,5 for (g); * indicates P<0.05, ** indicates P<0.01, two-way ANOVA (e), one-way ANOVA (g), and post-hoc Bonferroni’s test; Data in (e) and (g) are shown as means ± SEM.

To examine the role of IGF1 in hMGE development, we treated 6-week-old MGEOs, which were transplanted with or without iMG at 4 weeks of age, with either a carrier control (PBS), recombinant human IGF1 proteins (IGF1), or IGF1-neutralizing antibodies (anti-IGF1) for 48 hours. BrdU was added during the last 4 hours of the 48-hour treatment for precise evaluation of proliferation cells (Fig. 5a). We observed that the presence of iMG increased the density of BrdU^+^NKX2.1^+^ cells in MGEOs (Fig. 5d, e). More importantly, IGF1 treatment drastically increased the density of BrdU^+^NKX2.1^+^ cells in MGEOs without iMG transplantation, whereas IGF1-neutralizing antibodies abolished the elevated density of BrdU^+^NKX2.1^+^ cells in organoids with iMG (Fig. 5d, e). On the other hand, the densities of iMG in MGEOs were not different among the various treatments (Fig. 5d and Extended Data 13).

To definitively demonstrate that the IGF1 is the key regulator mediating microglial regulation of interneuron development, we generated IGF1 loss-of-function mutation (IGF1 KO) hESC lines using CRISPR/Cas9-based non-homology end jointing (Extended Data Fig. 14). We then generated MGEOs transplanted with no iMG, control iMG, or IGF1 KO iMG at 4 weeks of age and examined the MGE progenitor proliferation by quantifying the density of BrdU and NKX2.1 double-positive cells at 6 weeks old (Fig. 5a). We observed that the MGE progenitor proliferation increased in MGEOs transplanted with control iMG but not in those transplanted with IGF1 KO iMG (Fig. 5f, g). Addition of recombinant human IGF1 rescued the phenotype associated with IGF1 KO iMG (Fig. 5f, g). In summary, our results supported that iMG exert their function in promoting MGE progenitor proliferation through IGF1.

## Discussion

In summary, our findings provide critical insight into the role of microglia in the development of hMGE and their influence on GABAergic interneurons (Fig. 5h). We demonstrate that microglia exhibit a region-specific distribution pattern, closely interacting with proliferating progenitors in the hMGE. Further studies based on transcriptomic analysis and 3D MGE neuroimmune organoid models demonstrated a clear function of microglia in MGE progenitor proliferation through IGF1 pathway. Our study supports that microglia contribute to sustained neurogenesis in the hMGE.

Studies from rodents have showed region-specific interaction between radial glia and microglia. For example, microglia wrap their processes around radial glia projections in the embryonic mouse hypothalamus and respond to immune challenges ^41^. Radial glia endfeet contact with meningeal microglial precursors and regulate microglia development through integrin αVβ8-TGFβ1 signaling in the embryonic mouse cortex ^42^. Our findings support that microglia closely interact with radial glia in the SVZ of hMGE (Fig. 1l). Furthermore, our data with neuroimmune organoid models indicate that microglia enhance the proliferation of radial glia by secreting IGF1 (Fig. 4 and Fig. 5), highlighting a vital neuroimmune mechanism in brain development. Further studies are needed to understand the mechanism underlying the intimate and preferential association of radial glia and microglia in hMGE.

Brain organoids have been used to model various aspects of brain development and disease *in vitro* ^43–50^. This *in vitro* model enables mechanistic studies of human-specific aspects of brain development. We adapted a ventral telencephalon-oriented organoid model and demonstrated sequential presence of MGE progenitors, hMGE unique DEN-like organization, and MGE-derived PV^+^ and SST^+^ interneurons, resembling the developmental trajectory of the developing hMGE (Fig. 3b-g). One limitation of brain organoid models is the lack of endogenous microglia and the short survival duration of transplanted microglia. Here, we show that the presence of trophic factors, recombinant human CSF1, IL-34, and TGFβ, support iMG survival up to 12 weeks post transplantation. Importantly, those iMG resemble human primary microglia morphologies, transcriptomics, (Fig. 3h-n), and physical proximity to proliferating radial glia and progenitors (Fig. 3o-q). Thus, our newly established neuroimmune MGEOs serve as a reliable model to study microglial regulation of hMGE development.

Our study results have significant implications for understanding the development of GABAergic interneurons in human brains and the potential etiology of neurological and psychiatric disorders associated with microglia and interneuron dysfunction. Epidemiological studies have shown lower levels of IGF-1 in cerebrospinal fluid of children with autism ^51,52^. Additional human and mouse studies have demonstrated the pottial efficacy of IGF1 administration in ameliorating autistic features ^53–57^. Our study indicates that microglia are the main source of IGF1 in the embryonic human brain (Fig. 2e-i, and Extended Data Fig. 6a, b) and that IGF1 promotes interneuron production (Fig. 5), suggesting the possibility that low levels of IGF1 could result in interneuron deficits.

## Methods

### Human Tissue Samples

De-identified human specimens were collected from the Autopsy Service in the Department of Pathology at the University of California San Francisco (UCSF) (Table S1), with previous patient consent in strict observance of the legal and institutional ethical regulations. The autopsy consents and all protocols for human prenatal brain tissue procurement were approved by the Human Gamete, Embryo and Stem Cell Research Committee (Institutional Review Board GESCR# 10-02693) at UCSF. All specimens received diagnostic evaluations by a board-certified neuropathologist as control samples and were free of brain-related diseases. The diagnostic panel included the assessments of neural progenitor and immune cells using immunohistochemistry (IHC) to ensure that all control cases were not affected by any inflammatory diseases. Tissues used for snRNA-seq were snap-frozen either on a cold plate placed on a slab of dry ice or in isopentane on dry ice. Tissues later used for immunohistochemistry (IHC) were cut coronally into 1 mm tissue blocks, fixed with 4% paraformaldehyde for two days, cryoprotected in a 15-30% sucrose gradient, embedded in optimal cutting temperature (OCT) compound, and sectioned at 30 μm with a Leica cryostat and mounted onto glass slides.

### Authentication of cell lines used

All human induced pluripotent stem cell (iPSC) and human embryonic stem cell (hESC) lines used in this work have been karyotyped and regularly tested for mycoplasma. The eWT-1323-4 hiPSC line ^45^ (female, RRID: CVCL_0G84) was obtained from the Conklin Laboratory (University of California, San Francisco (UCSF). WA09/H9 (female, RRID: CVCL_9773, NIH registration number: NIHhESC-10_0062) and WA01/H1 (male, RRID: CVCL_9771, NIH registration number: NIHhESC-10-0043) were obtained from the WiCell Research Institute (Madison, WI, USA).

### Mouse

All mice were handled according to the guidelines of the Institutional Animal Care and Use Committee at the University of California, San Francisco. Wild type C57/B6 mice were purchased originally from Taconic Biosciences and bred in the laboratory. Males and females were mated together for one night for timed pregnancy. The noon after the mating night was noted as embryonic day (E) 0.5. In the EdU labeling experiment, a single dose of EdU (10 mg/kg; provided in Click-it EdU Alexa Fluor 647 Imaging Kit from Invitrogen, #C10340) was injected intraperitoneally into pregnant mice at E14.5. At E14.5 and E16.5, the pregnant dams were sacrificed, and the fetal brains were collected, fixed in 4% PFA at 4°C overnight, cryo-preserved in 30% sucrose at 4°C overnight, then cryo-sectioned at 40 μm with a Leica cryostat

### Human Pluripotent Stem Cell (hPSC)-Derived MGE Organoids (MGEO)

hPSC-derived MGE organoids were generated largely following the previous established protocol ^35,36^. Briefly, 1323-4 hiPSCs or WA01/H1 and WA09/H9 hESCs were expanded in StemFlex Basal Medium (Gibco, A3349401). After reaching 80% coverage, hPSCs cultured on Matrigel were dissociated into clumps using ReLesR (Stemcell Technologies, 100-0483) and equally distributed into a V-bottom 96-well ultra-low attachment PrimeSurface® plate (S-bio, MS-9096VZ). The Rho Kinase Inhibitor Y-27632 (10 μM) was added during the first 24 hours of neural induction to promote survival. Between days 0-5, organoids were cultured in neural induction media (DEME/F12 medium, 20% knockout serum, 1% NEAA,0.5% Glutamax, and 0.1mM β-ME, and 1% penicillin-streptomycin)) supplemented with the SMAD inhibitors SB431542 (10 μM) and dorsomorphin (5 μM). Between days 6-24, organoids were cultured in neural differentiation media (Neurobasal A medium, 2% B27 supplement, 1% GlutaMAX, and 1% penicilin-streptomycin) supplemented with human recombinant EGF (20ng/ml) and human recombinant FGF2 (20ng/ml). Between days 25-43, organoids were maintained in neural differentiation media supplemented with human recombinant brain-derived neurotrophic factor (BDNF) (20 ng/ml) and human recombinant neurotrophin 3 (NT3) (20 ng/ml). The media were also supplemented with 5 μM wnt-inhibitor IWP-2 during days 4-23; 100 nM smoothened agonist (SAG) during days 12-23, 100 nM retinoic acid during days 12-15, and 100 nM allopregnanole during days 16-23 for ventral forebrain patterning. Each organoid was induced and generated in a single 96-well and would be moved 6-well plates for long-term culture after week 5. All the media and supplements used for organoids culture were the same as the previous published protocol^35,36^.

### Induced microglia (iMG)

Induced microglia cells were generated from WA01/H1 or WA09/H9 hESC cells under defined conditions as previously described ^37^. Briefly, hESCs were differentiated into CD43-expressing hematopoietic progenitor cells (HPCs) for 12 days using the STEMdiff™ Hematopoietic Kit (Stemcell, 05310). HPCs were differentiated for 24 days using the STEMdiff™ Microglia Differentiation Kit (Stemcell, 100-0019) and matured for an additional 4 days using the STEMdiff™ Microglia Maturation Kit (Stemcell, 100-0020) before being added to organoid cultures for co-culture.

### iMG-organoid engraftment and co-culture

Mature iMG were immediately added to 4-week-old MGE organoids in 96-well ultra-low attachment PrimeSurface® plates at 80-100 ×10^3^ microglia/organoid. Trophic factors (100 ng/mL IL34 (Peprotech, 200-34), 25 ng/mL CSF1(Peprotech, 300-25), 50 ng/mL TGFβ1(Peprotech, 100-21)) were added to the culture medium to support microglial survival. One week post transplantation, co-cultured organoid-microglia (neuroimmune organoids) were transferred to a 6-well plate and placed on an orbital shaker. The co-cultures were then maintained following the usual organoid maintenance protocol in addition with trophic factors.

### Pharmacological manipulation of MGEOs

MGEOs were treated with PBS, 100 ng/mL recombinant human IGF1 (Abcam, ab269169) or 1 μg/mL IGF1-neutralizing antibodies (Abcam, ab9572) for 48 hours. 10 μΜ BrdU (Abcam, ab142567) was added during the last 4 hours to label proliferating cells. Organoids were then collected right after treatment.

### Immunohistochemistry (IHC)

We followed the IHC protocol as previously reported ^12,30^. Fixed human samples were fixed and cryo-sectioned as described above in **Human Tissue Samples**. Mouse samples were prepared as described above in **Mouse**. Organoids were fixed in 4% paraformaldehyde for 30-45 mins at the room temperature and then cryopreserved in 30% sucrose in PBS overnight. The organoids were then embedded in OCT (SCIgen, #4586) and cryo-sectioned at 14 μm with a Leica cryostat.

Mounted human slides were defrosted overnight at 4°C and then dried at 37°C for 3 hours. Mounted organoid and mouse slides were directly dried at 37°C for 30 mins. Antigen retrieval was performed for 10-12 min at 95-99°C with antigen retrieval buffer (BD Pharmingen, #550524). After antigen retrieval, tissue slices were washed with 1x PBS plus 0.1 % Triton X-100, then blocked in blocking buffer (5% serum, 1% bovine serum albumin (BSA), and 0.1% Triton X-100 in PBS) for 1.5 hours at room temperature before proceeding to incubation with primary antibodies overnight at 4°C. After washing, sections were incubated with species-specific secondary antibodies conjugated to Alexa Fluor dyes (1:500, Invitrogen) for 2 hours at room temperature. For human slides, TrueBlack Lipofuscin Autofluorescence Quencher (1:20 in 70% alcohol, Biotium, #23007) was applied for 3 minutes to block autofluorescence. For EdU staining, EdU working solution was applied to E16.5 mouse brain slices after secondary antibodies application following the manufacturer’s instructions. Nuclei were counterstained with DAPI (1:1000 from 1 mg/ml stock, Invitrogen, 2031179) for five minutes. Images were captured using a Leica Stellaris 8 confocal microscope. For organoid experiment, three slices of each organoid were imaged, quantified and averaged for the final statistical analysis.

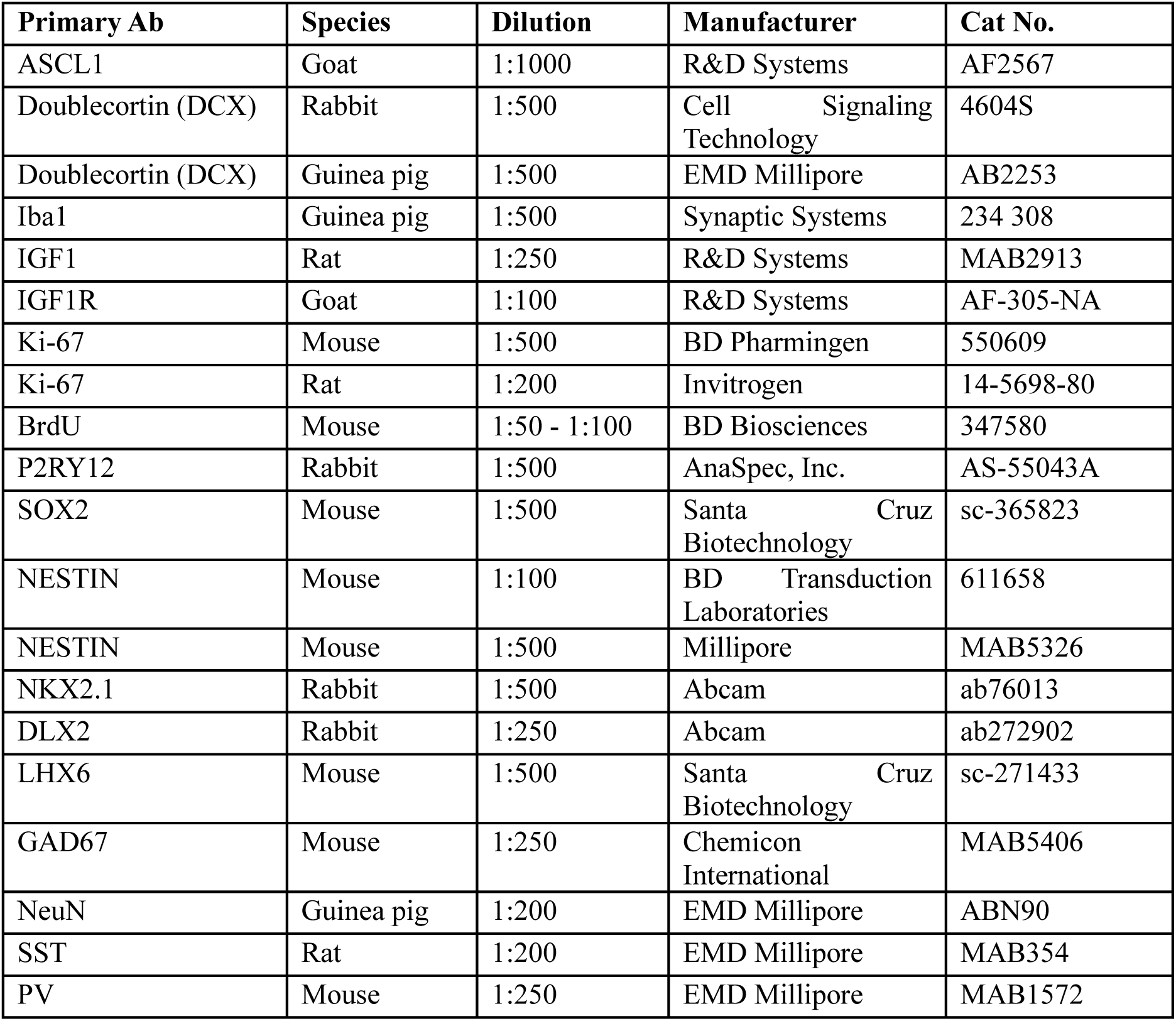

| Primary Ab | Species | Dilution | Manufacturer | Cat No. |
| --- | --- | --- | --- | --- |
| ASCL1 | Goat | 1:1000 | R&D Systems | AF2567 |
| Doublecortin (DCX) | Rabbit | 1:500 | Cell Signaling Technology | 4604S |
| Doublecortin (DCX) | Guinea pig | 1:500 | EMD Millipore | AB2253 |
| Iba1 | Guinea pig | 1:500 | Synaptic Systems | 234 308 |
| IGF1 | Rat | 1:250 | R&D Systems | MAB2913 |
| IGF1R | Goat | 1:100 | R&D Systems | AF-305-NA |
| Ki-67 | Mouse | 1:500 | BD Pharmingen | 550609 |
| Ki-67 | Rat | 1:200 | Invitrogen | 14-5698-80 |
| BrdU | Mouse | 1:50 - 1:100 | BD Biosciences | 347580 |
| P2RY12 | Rabbit | 1:500 | AnaSpec, Inc. | AS-55043A |
| SOX2 | Mouse | 1:500 | Santa Cruz Biotechnology | sc-365823 |
| NESTIN | Mouse | 1:100 | BD Transduction Laboratories | 611658 |
| NESTIN | Mouse | 1:500 | Millipore | MAB5326 |
| NKX2.1 | Rabbit | 1:500 | Abcam | ab76013 |
| DLX2 | Rabbit | 1:250 | Abcam | ab272902 |
| LHX6 | Mouse | 1:500 | Santa Cruz Biotechnology | sc-271433 |
| GAD67 | Mouse | 1:250 | Chemicon International | MAB5406 |
| NeuN | Guinea pig | 1:200 | EMD Millipore | ABN90 |
| SST | Rat | 1:200 | EMD Millipore | MAB354 |
| PV | Mouse | 1:250 | EMD Millipore | MAB1572 |

### 3D Reconstruction and Image Analysis

Three-dimensional (3D) reconstructions were generated using Imaris software (Oxford Instrument). For distance analysis, microglia were reconstructed using the surface modules, while Ki-67^+^ cells or DAPI^+^ cells were reconstructed with spot modules. The distance of the center of each cell (spot) to the nearest microglia (surface) was determined unbiasedly by Imaris. The distance distribution of Ki-67^+^ and Ki-67^−^ cells to the nearest microglia were calculated accordingly.

### Single nucleus preparation

Single-nucleus suspensions were prepared from postmortem human samples. About 50 mg of sectioned freshly frozen human brain tissue was homogenized in lysis buffer (0.32 M sucrose, 5 mM CaCl_2_, 3 mM MgAc_2_, 0.1 mM EDTA, 10 mM Tris-HCl, 1 mM Dithiothreitol, 0.1% Triton X-100 in DEPC-treated water) plus 0.4 U/μL RNase inhibitor (Takara, Cat #2313A) on ice. Then, the homogenate was loaded into a 30 mL thick polycarbonate ultracentrifuge tube (Beckman Coulter, Cat # 355631), and 9 mL of sucrose cushion solution (1.8 M sucrose, 3 mM MgAc_2_, 1 mM Dithiothreitol, 10 mM Tris-HCl in DEPC-treated water) was added to the bottom of the tube. The tubes with tissue homogenate and sucrose cushions were then ultra-centrifuged at 107,000 xg for 2.5 hours at 4°C. The pellet was recovered in 250 uL ice-cold PBS for 20 min, resuspended in nuclei sorting buffer (PBS, 1%BSA, 0.5 mM EDTA, 0.1U/μL RNase inhibitor) and filtered through a 40 μm cell strainer to get single nucleus suspensions for the following fluorescence-activated cell/nucleus sorting.

### Single cell preparation

Single cell suspensions of 1323-4 hiPSC-derived organoids were prepared with neural tissue dissociation kits (P) (Miltenyi Biotec, 130-092-628) following manufactural instructions. Briefly, 12-16 organoids per experimental condition were processed through a gentle two-step enzymatic dissociation procedure as instructed. 5 mg/ml Actinmycin D (Sigma-Aldrich, A1410), 10 mg/ml Anisomycin (Sigma-Aldrich, A9789), and 10 mM Triptolide (Sigma-Aldrich, T3652) was added additionally prior to the tissue digestion to inhibit the cellular transcriptome. Following digestion, organoids were mechanically triturated using fire-polished glass pipettes, filtered through a 40 μm cell strainer test tube (Corning, 352235), pelleted at 300 x g for 5 minutes, and washed twice with DBPS before proceeding to 10x Genomics scRNA library preparation. For samples that need fluorescence-activated cell sorting, the single cell pellet was resuspended in cell sorting buffer (DPBS, 1% BSA, 0.1 U/μL RNase inhibitor).

### Fluorescence-activated cell/nucleus sorting (FACS/FANS)

Single nuclei suspensions from fresh-frozen human samples was stained with antibodies of PU.1 (Cell Signaling Technology, 81886S, 1:100), and OLIG2 (Abcam, ab225100, 1:2500) overnight at 4°C. PU.1 and OLIG2 antibodies were conjugated with fluorescence upon purchase. DAPI (1:1000) was added for 5 min on the second day. Single cell suspensions from organoids were stained with DAPI (1:1000) for 5 min in cell sorting buffer (DPBS, 1% BSA, 0.1 U/μL RNase inhibitor). The single nucleus/cell suspension was then centrifuged at 300 x g for 5 min, resuspended in nuclei/cell sorting buffer, and filtered through a 40 μm cell strainer for final analysis and sorting through a FACS Aria II cell sorter (BD Biosciences). Target cells were collected in nucleus/cell sorting buffer for future sequencing library preparation.

### Single cell/nucleus- RNA library preparation

Nuclei/cells were counted using a hemocytometer and resuspend to a final concentration of 300-1,000 cells/nuclei per μL in PBS. Single nuclei/cell RNA-seq libraries were prepared using 10x Genomics Chromium Next GEM single cell v3.1 kit according to the manufacturer’s instruction, targeting for 5,000 nuclei/cells per sample. Single cell/nucleus libraries were then sequenced on the NovaSeq 6000 machine, aiming the sequencing depth of 50,000 reads per cell.

### Single cell/nucleus-RNA-seq data analysis

Sequencing results were then aligned to the GRCh38 genome (gex-GRCh38-2020-A) using CellRanger v6.1.2 (10x Genomics). --include-introns was used to include pre-mature mRNA in single-nucleus samples. Gene counts then underwent a doublet removal step using DoubltFinder v2.0.3 (https://www.cell.com/cell-systems/fulltext/S2405-4712(19)30073-0). Nuclei/cells with detected genes ranged from 500-10000, minimum of 1000 UMI, and less than 5% reads mapped to mitochondria genes were kept.

The output (count matrix) was used as the main input file for all downstream analysis using Seurat v 5.1.0. Cells/nuclei that had a UMI of less than 1,000 or more than 100,000 were filtered out. For snRNA-seq, mitochondrial counts more than 2% were also filtered out. MALAT1, mitochondrial genes (MT-), ribosomal protein encoding genes (RPS- and RPL-), and hemoglobin genes (HB-) were excluded for further analysis. Standard data normalization, variable features identifications, linear transformations, dimensional reduction, UMAP embedding, and unsupervised clustering were conducted using the standard Seurat pipeline^33^. The cell type cluster identification was determined based on expression of known marker genes, as is shown in fig. S1, S6, and S8. For scRNA-seq of GFP-labelled iMG, iMG were purified future *in silico* using canonical microglia/macrophage markers, including AIF1, CX3CR1, C3, PTPRC, ITGAM, and CD68.

Cell-cell interaction analysis was conducted using CellChat v2^34^. For development-based analysis, independent CellChat files were generated from ‘embryonic’ and ‘perinatal’ Seurat objects, and a comparison analysis was conducted between them. A heatmap was made via Graphpad Prism 9 according to the interacting probability of significant ligand-receptors interactions involved in microglial (MG) regulation of interneurons (CIN).

Differently expressed genes (DEGs) analysis was conducted based on the Seurat-default non-parametric Wilcoxon rank sum test. Pathways with enriched DEGs were generated using Enrichr (https://maayanlab.cloud/Enrichr/#) based on the Reactome Pathway Database, Kyoto Encyclopedia of Genes and Genomes (KEGG), Gene Expression Omnibus (GEO), and the Gene Ontology (GO) database.

### CRISPR/Cas9 Gene Editing

A WA09/H9 stem cell line with an IGF1 loss-of-function mutation (IGF1 KO) was generated using CRISPR/Cas9-based non-homology end joining largely following the protocol of the Alt-RΤΜ CRISPR-Cas9 System from Integrated DNA Technologies (IDT). The gRNA (5’TCGTGGATGAGTGCTGCTTC3’) is selected from Predesigned Alt-R CRISPR-Cas9 guide RNA (IDT). Equal amounts of crRNA and ATTO^TM550^ labelled tracrRNA (IDT, 1075927) were mixed to a final concentration of 100 μΜ, heated to 95°C for 5 min, and then cooled to room temperature to anneal followed by forming the RNP complex with Alt-RTM S.p. HiFi Cas9 Nuclease V3 (IDT,1081061) at room temperature for 20 min. The RNP complex was delivered to single-stem cell suspensions using the Neon^TM^ electroporation system (1400 V, 20 ms, 1 pulse) according to the manufacturer’s instructions. After electroporation, ATTO^TM550+^ cells were selected by FACS after three days of culture and sparsely seeded to form a single cell colony. A loss-of-function mutation cell line is selected by Sanger sequencing with out-of-frame mutations at the target site, followed by exclusion of any mutations at the top 5 potential off target sites. Further Sanger sequencing confirmation, RT-qPCR, and IHC were applied to confirm IGF1 KO.

### Data analysis and statistics

For all quantification, images were acquired blindly to genotype or treatment before quantification. Statistics were performed with Prism GraphPad (v10.1.0) as shown in each figure legend.

## Acknowledgments

We thank and honor the families who generously donated the tissue samples used in this study. We thank Dr. Alvarez-Buylla Arturo, Dr. Arnold Kriegstein, Dr. David H. Rowitch, and Dr. Stephen Fancy for discussion and thoughtful comments on the manuscript. We thank Mrejen Caroline of Weill Innovation Microscope Core and Center for Advanced Light Microscopy (CALM) core for technical supports in microscopy. Funding: National Institutes of Health/National Institute of Neurological Disorders and Stroke (NIH/NINDS) grant P01 NS083513 (MFP, EJH, XP), NIH/NINDS grant R01 NS094164 (XP), NIH/NINDS grant R01 NS108446 (XP), California Institute for Regenerative Medicine (CIRM) grant DISCO0-15949 (XP, TJN)NIH/NINDS grant R01 NS132595 (EJH),VA BLR&D Merit Review Award I01 BX001108 (EJH).

## Author contributions

D.Y. and X.P. conceived and designed the study. M.F.P., T.J.N., E.J.H., X. P. supervised the study. D.Y., A.W., B.Z., J.K., and J.J.C. performed histological and sequencing experiment with post-mortem human samples. D.Y., S.J., A.W., and S.F., performed experiments with organoids. D.Y., A.W., and S.J. conducted bioinformatic analysis. D.Y. and S.J. analyzed most results and made the figures. S.J., D.Y., and A.W. wrote methods. D.Y. and X.P. wrote the manuscript with comments from all authors.

## Data and materials availability

All data are available in the main text or the supplementary materials. Raw sequencing datasets, processed Seurat objects, and custom code used in the analyses were deposited to Gene Expression Ominibus (GEO), Zenodo, and Github. All other data are available upon request.

