## extended data for "Microglia regulate GABAergic neurogenesis in prenatal human brain through IGF1"

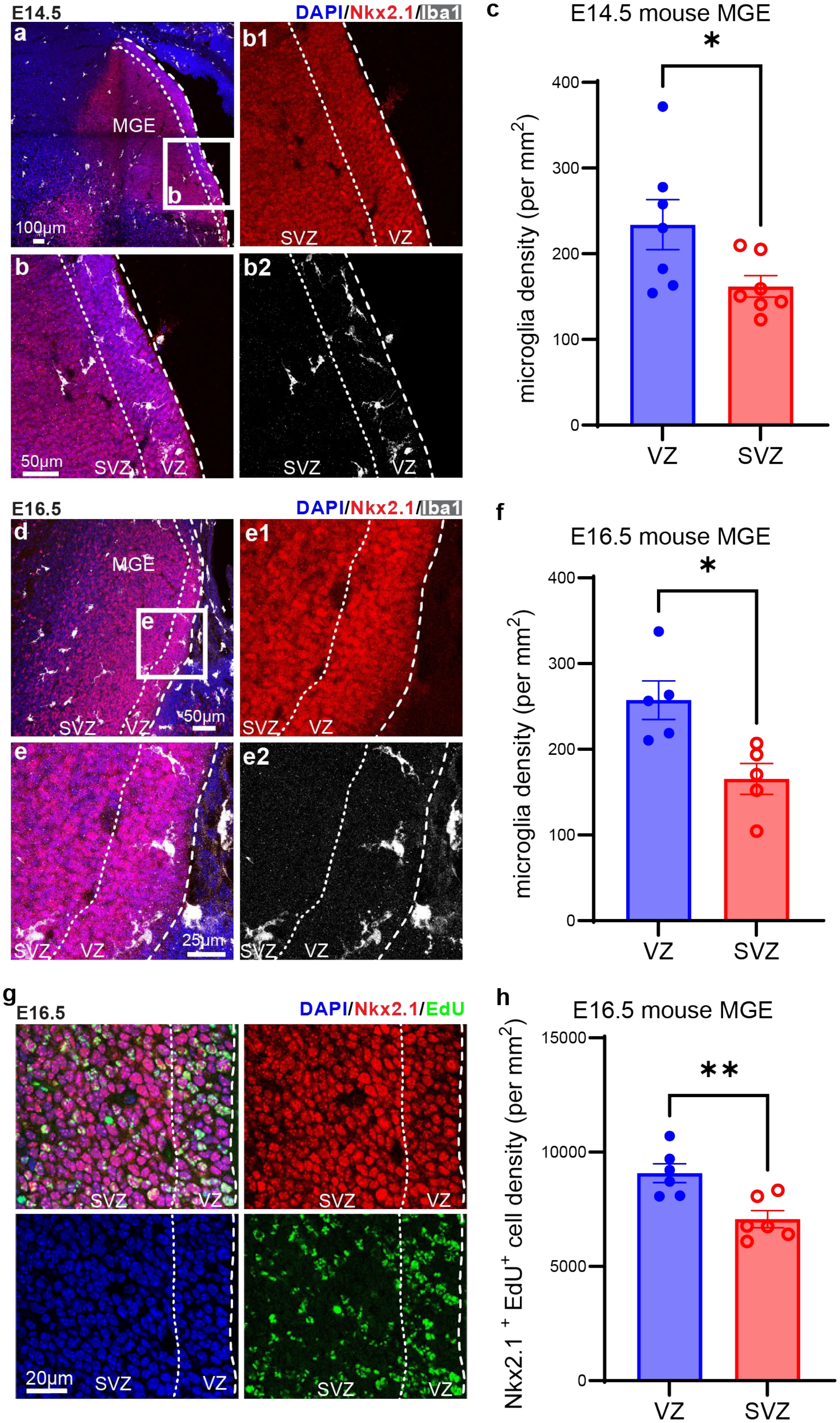


**Extended Data Fig. 1 | The distribution of microglia and proliferation progenitors in mouse MGE. a-b.** IHC images showing the distribution of Iba1$^+$ microglia in E14.5 mouse MGE. MGE were labelled by Nkx2.1. **c.** Bar graph showing the density of Iba1$^+$ microglia is significantly higher in the VZ of E14.5

mouse MGE. **d-e**. IHC images showing the distribution of Iba1$^+$ microglia in E16.5 mouse MGE. **f**. Bar graph showing the density of Iba1$^+$ microglia is significantly higher in the VZ of E16.5 mouse MGE. **g**. IHC images showing the distribution of EdU$^+$Nkx2.1$^+$ active progenitors in the E16.5 mouse MGE with EdU injected at E14.5. **h**. Bar graph showing the density of Nkx2.1$^+$EdU$^+$ active progenitors is significantly increased in the VZ of E16.5 mouse MGE with EdU injected at E14.5. *P<0.05, **P<0.01; N=7, 5, 6 for (c), (f), (h), respectively; unpaired t test. Data are shown as means ± SEM.

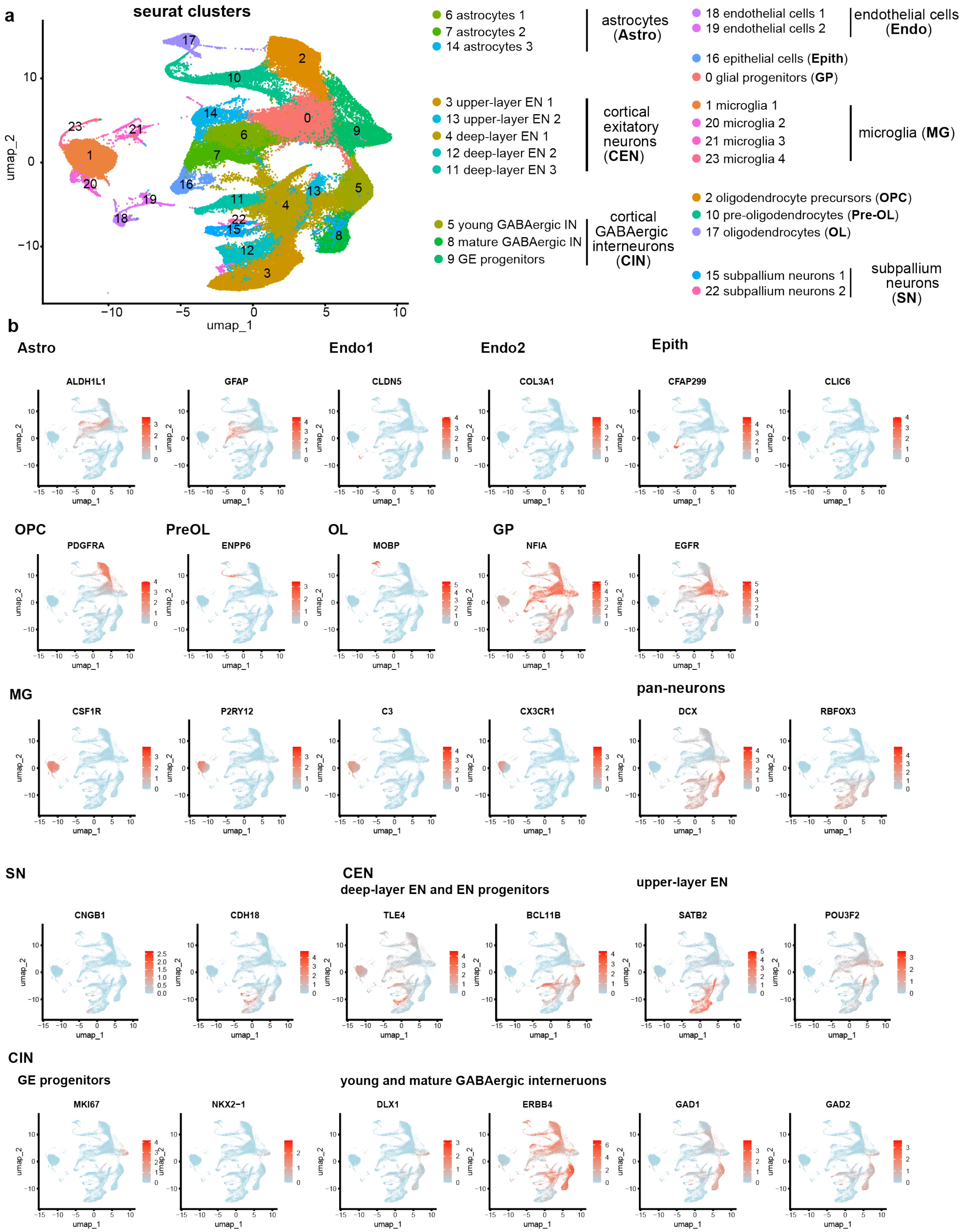


**Extended data Fig. 2 | Identification of cell types in human snRNAseq data. a**,.UMAP plots of Seurat clusters and cell type annotations. **b**, Feature plots showing selected canonical markers for different types of cells, including ALDH1L1 and GFAP for astrocytes (Astro), CLDN5 and COL3A for endothelial cells (Endo), CFAP299 and CLIC6 for epithelial cells (Epith), PDGFRA for oligodendrocyte precursor cells (OPC), ENPP6 for pre-myelinating oligodendrocytes (Pre-OL), MOBP for oligodendrocytes (OL), NFIA and EGFR for glia progenitors (GP), CSF1R, P2RY12, C3, and CX3CR1 for microglia (MG), DCX and RBFOX3 for pan-neurons, CNGB1 and CDH18 for subpallium neurons (SN), TLE4, BCL11B, SATB2 and POU3F2 for cortical excitatory neurons (CEN), MKI67 and NKX2-1 for GE progenitors, DLX1, LHX6, GAD1, and GAD2 for GABAergic interneurons. GE progenitors (cluster 9, NKX2.1$^{+}$MKI67$^{+}$), young GABAergic interneuron (cluster 5, DCX$^{+}$RBFOX3$^{low}$GAD1$^{+}$ GAD2$^{+}$), and mature GABAergic interneurons (RBFOX3$^{+}$DCX$^{low}$GAD1$^{+}$ GAD2$^{+}$) were combined as cortical interneurons (CIN) for the following analysis.

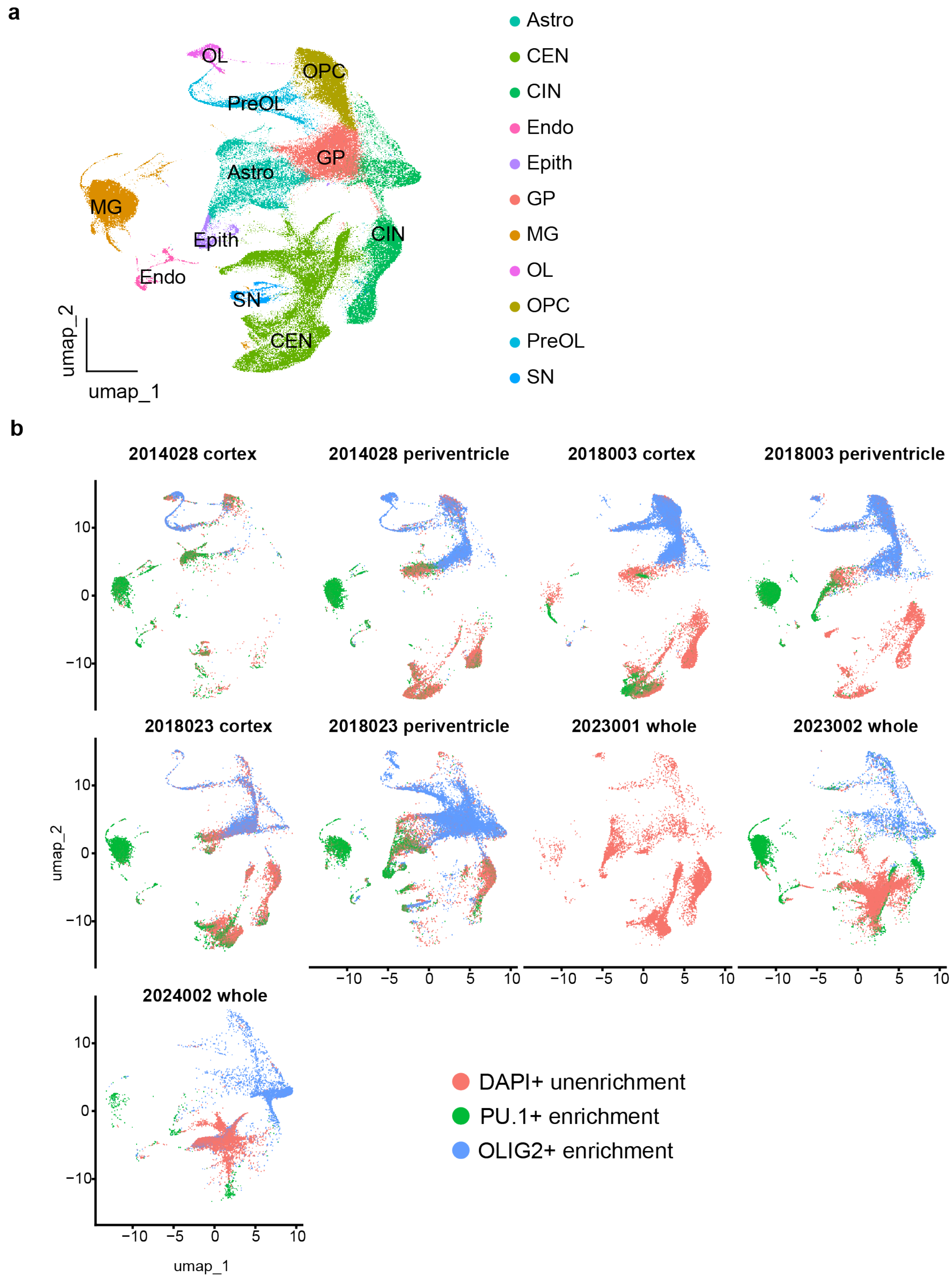


**Extended data Fig. 3 | UMAP plots showing cell clusters composition for different enrichment strategies from each sample. a**, UMAP plots with cell types of the clusters annotated. **b**, UMAP plots of each sample according to enrichment strategies.

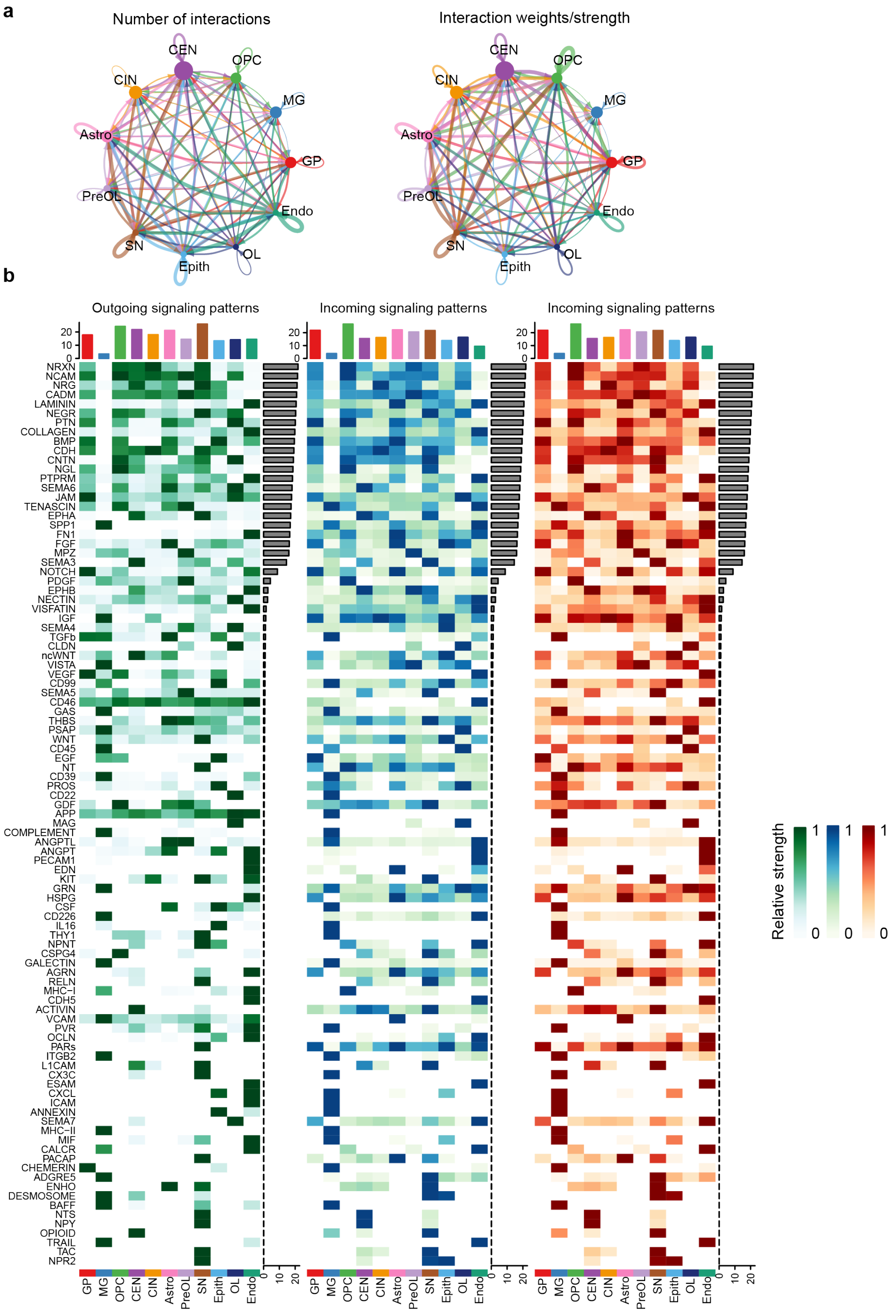


**Extended Data Fig. 4 | Cell-cell communication pathways revealed by CellChat analysis. a**, Circle plots showing the number of interactions and interaction weights/strength among different types of cells. **b**, Heatmaps showing the significant signaling pathways in terms of incoming, outgoing, and overall signaling.

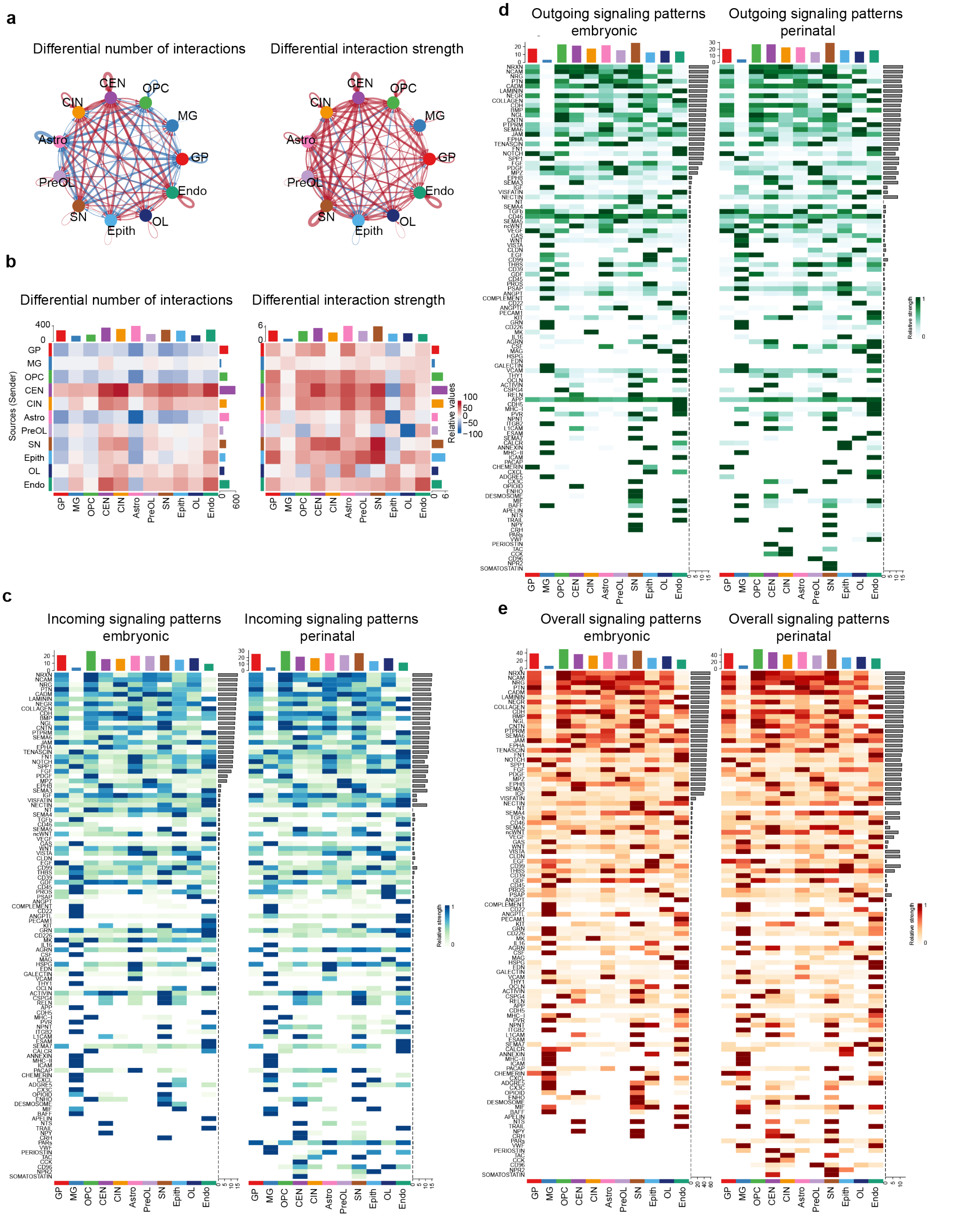


**Extended Data Fig. 5 | Differential cell-cell communication pathways at embryonic and perinatal stages. a**, Circle plots and **b**, heatmaps showing the number and weights/strength of differential interactions between embryonic and perinatal stages. **c**-**e**, Heatmaps showing signaling pathways in each cell type at embryonic and perinatal stages in terms of incoming (c), outgoing (d), and overall signaling (e).

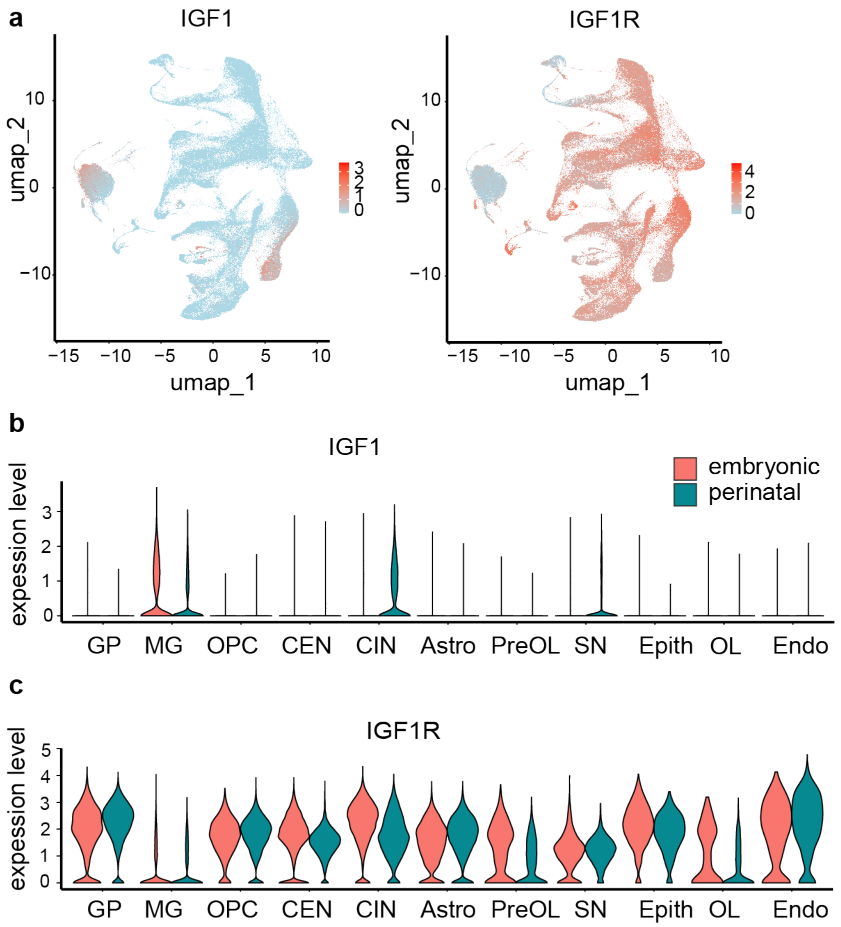


**Extended data Fig. 6 | Feature plots (a) and violin plots (b-c) showing the expression pattern of IGF1 and IGF1R.**

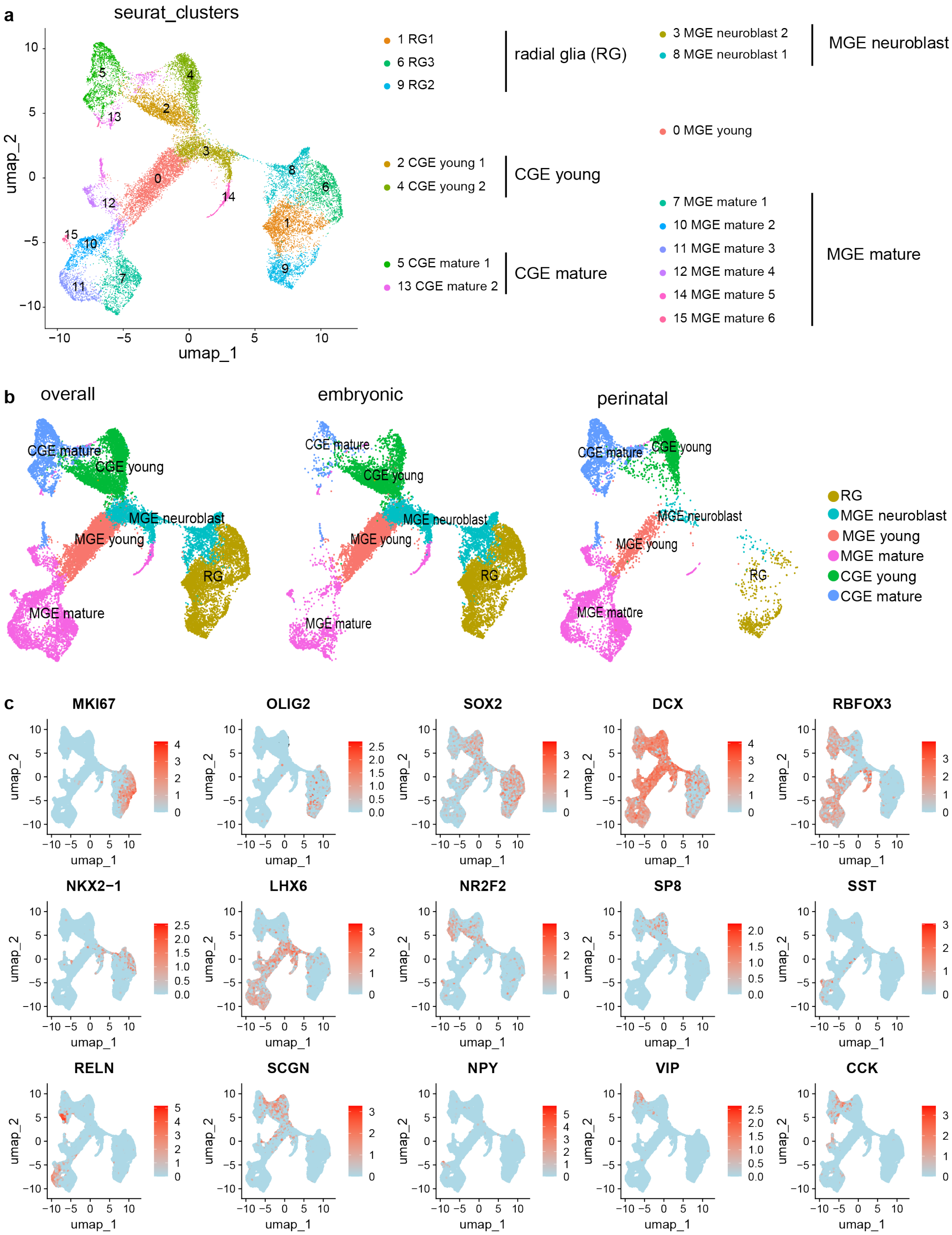


**Extended data Fig. 7 | Identification of interneuron subtypes in human snRNAseq data. a**, Interneuron subtype recognition according to Seurat clustering and canonical markers. **b**, UMAP of interneuron subclusters in the embryonic and perinatal stages. **c**, Feature plots of canonical markers. Radial glia (RG) are cell clusters that express high MKI67, OLIG2, and SOX2; MGE neuroblasts are cell clusters with residual MKI67, SOX2 and NKX2-1 expression and starting to express DCX and LHX6, corresponding to the neuroblast cells within DENs in the hMGE. MGE-derived young interneurons (MGE young) are cell clusters that do not express MKI67 and NKX2-1, but express high levels of DCX and LHX6 and low levels of RBFOX3. MGE-derived mature interneurons (MGE mature) are cell clusters that express high levels of RBFOX3 and LHX6 and low levels of DCX. CGE-derived young interneurons (CGE young) are cell clusters that do not express MKI67, but express high levels of DCX, NR2F2, and SP8, and low levels of RBFOX3. CGE-derived mature interneurons (CGE mature) are cell clusters that express high levels of RBFOX3, NR2F2, and LHX6, and low levels of DCX.

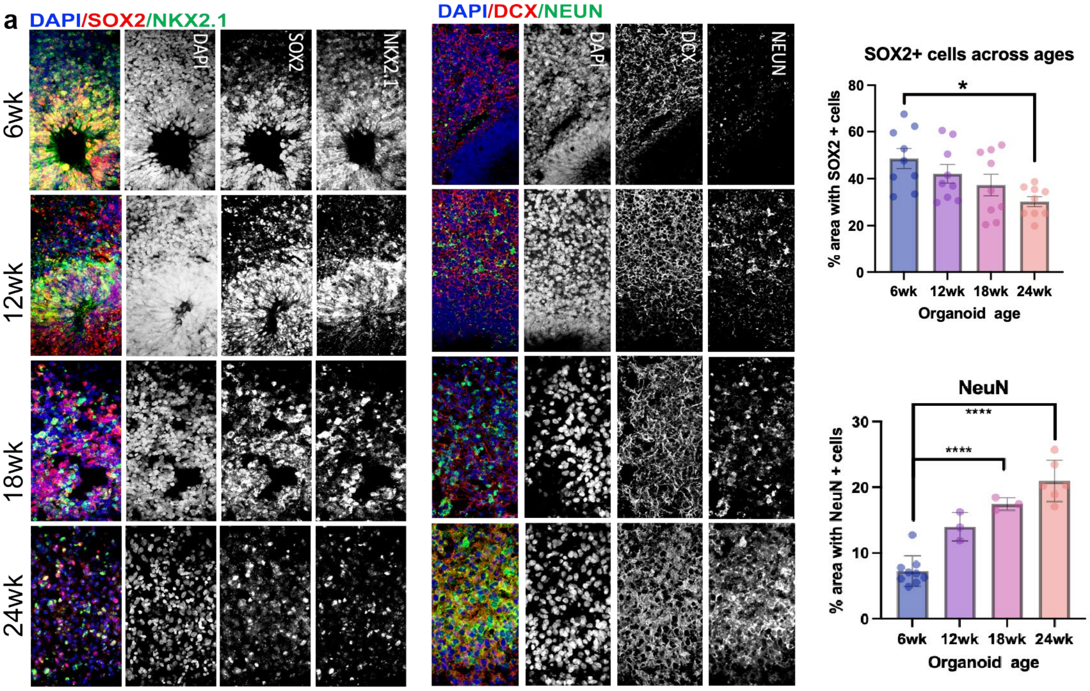


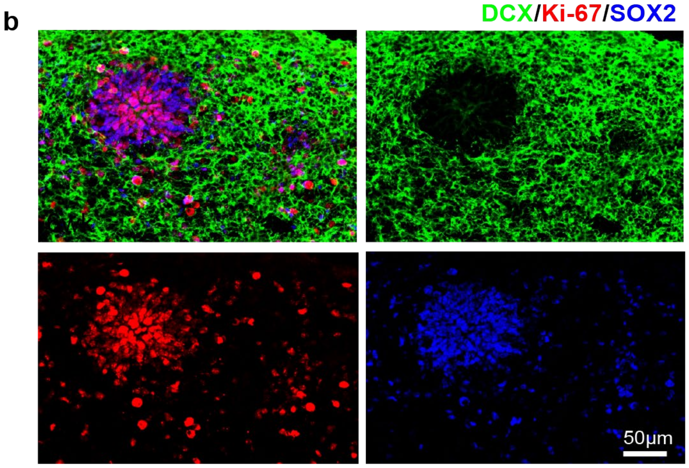


**Extended data Fig. 8 | Characterization of MGEOs.** **a**, MGEOs sequentially express markers for radial glia (SOX2) and postmitotic interneurons (DCX and NeuN) as they mature. $SOX2^{+}$ cells are high at 6 weeks-old and gradually decrease, whereas $NeuN^{+}$ neurons gradually increase as MGEOs age. **b**, Ki-$67^{+}$ cells seen in both $SOX^{+}DCX^{-}$ radial glia and $SOX2^{+}DCX^{+}$ neuroblasts in 6-week-old MGEOs.

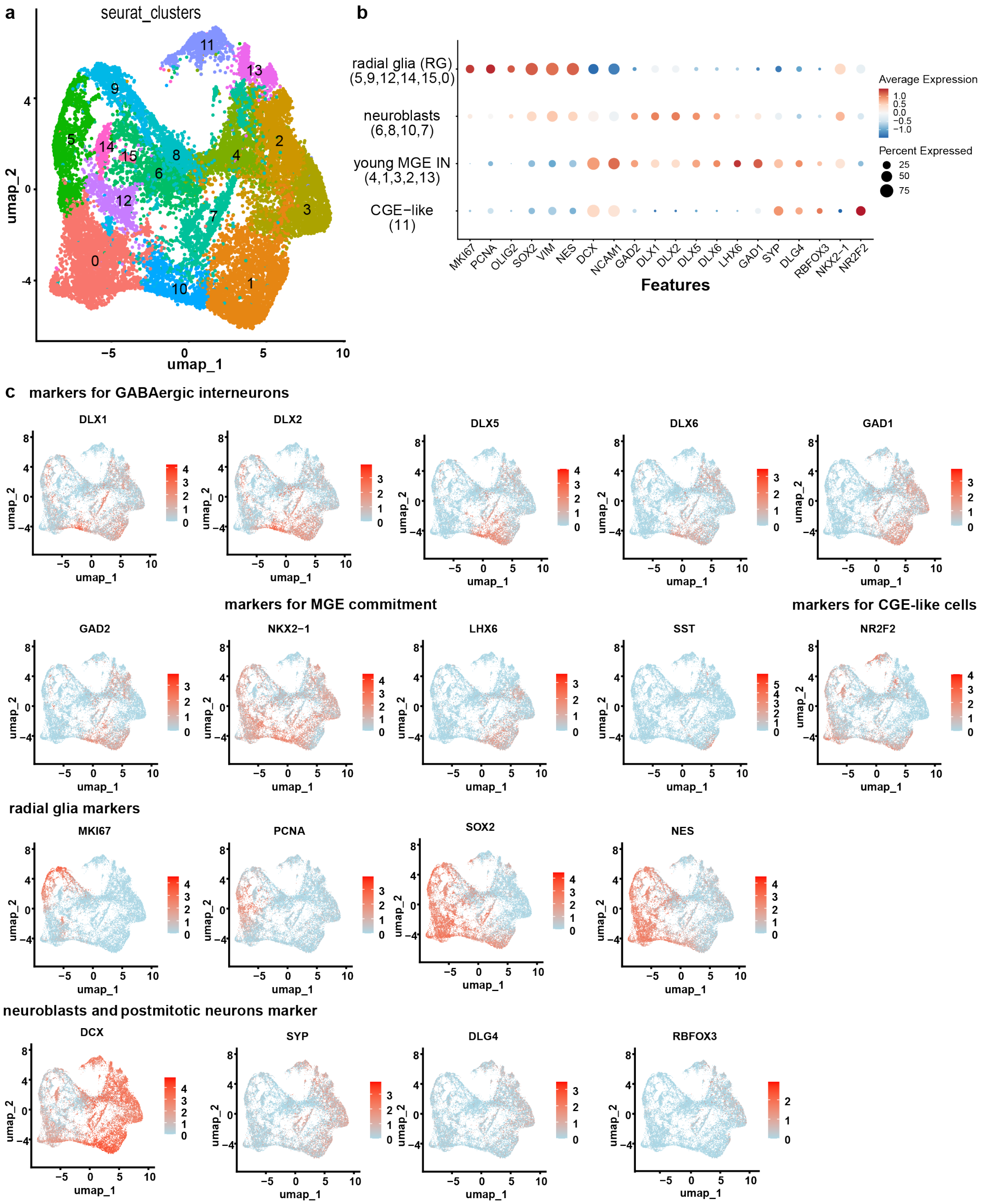


**Extended data Fig. 9 | Identification of cell types in 6-week-old MGEOs. a**, UMAP plots of unbiased clustering of MGEOs. **b**, Dot plots showing the level and cell percentage of canonical marker expression. **c**, Feature plots showing the expression of canonical markers. Radial glia (RG) are recognized as cell clusters that express high MKI67, PCNA, OLIG2, SOX2, VIM, and NESTIN (NES); neuroblasts are recognized as transiting clusters expressing both progenitor markers such as MKI67, PCNA, SOX2, and NES, as well as the young postmitotic neuron marker

DCX. Young MGE-derived GABAergic interneurons (Young MGE IN) are recognized as clusters that express DCX, NCAM, GAD1, GAD2, GAD5, GAD6, and low levels of SYP, DLG4, and RBFOX3.

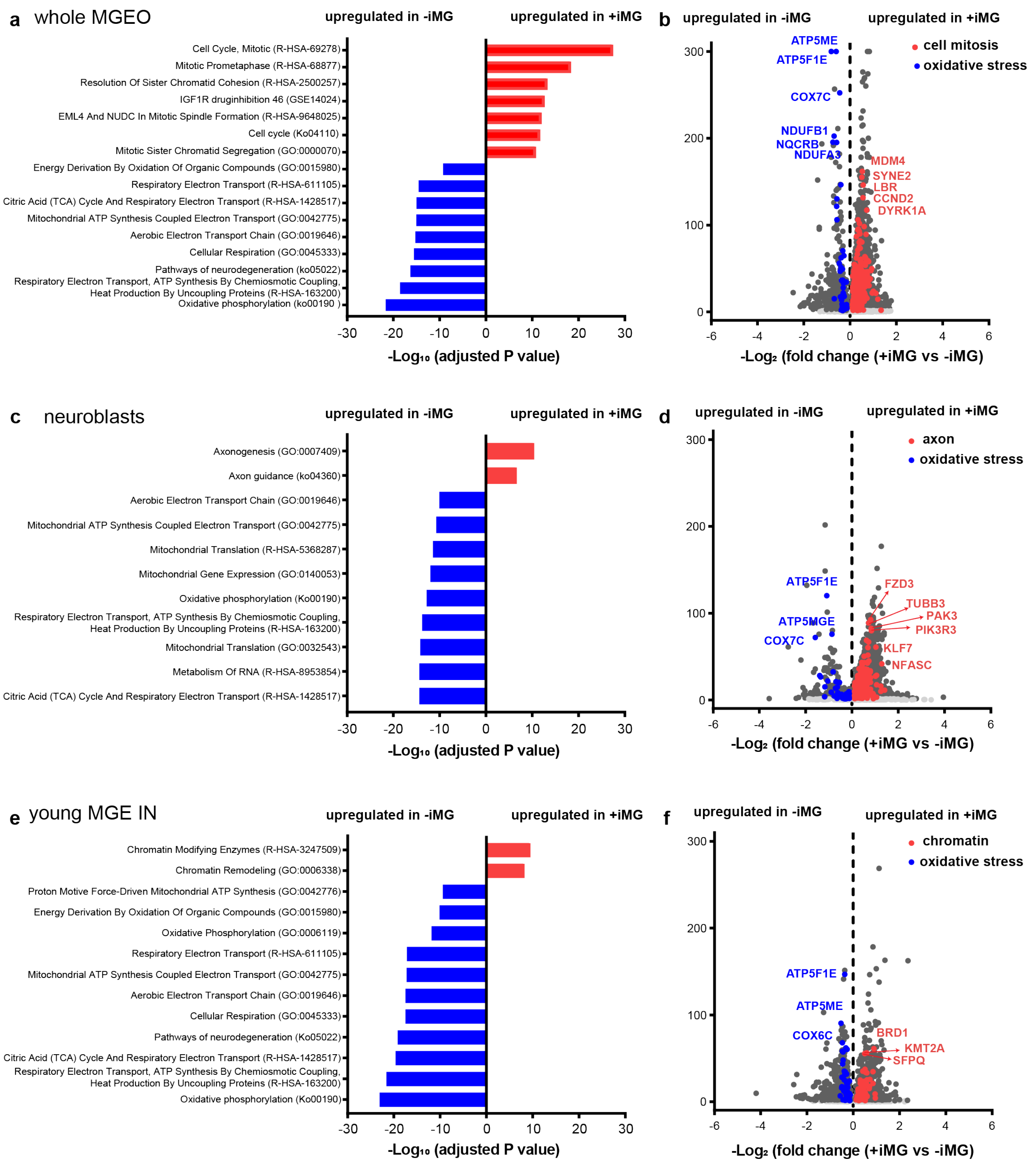


**Extended data Fig. 10 | DEGs and pathway analysis. a-b**, Pathway enrichment analysis plot (a) and volcano plot (b) of whole MGEO showing a higher expression of genes related to cell mitosis and lower expression of genes related to oxidative stress in MGEOs with iMG. **c-d**, Pathway enrichment analysis plot (c) and volcano plot (d) of neuroblasts cluster revealing an increased expression of genes related to axon development and decreased expression of genes related to oxidative stress in MGEOs with iMG. **e-f**, Pathway enrichment analysis plot (e) and volcano plot (f) of young MGE-derived GABAergic interneuron (young MGE IN) populations showing the presence of iMG leads to an increased the expression of genes related to chromatin metabolism and decreased expression of genes related to oxidative stress.

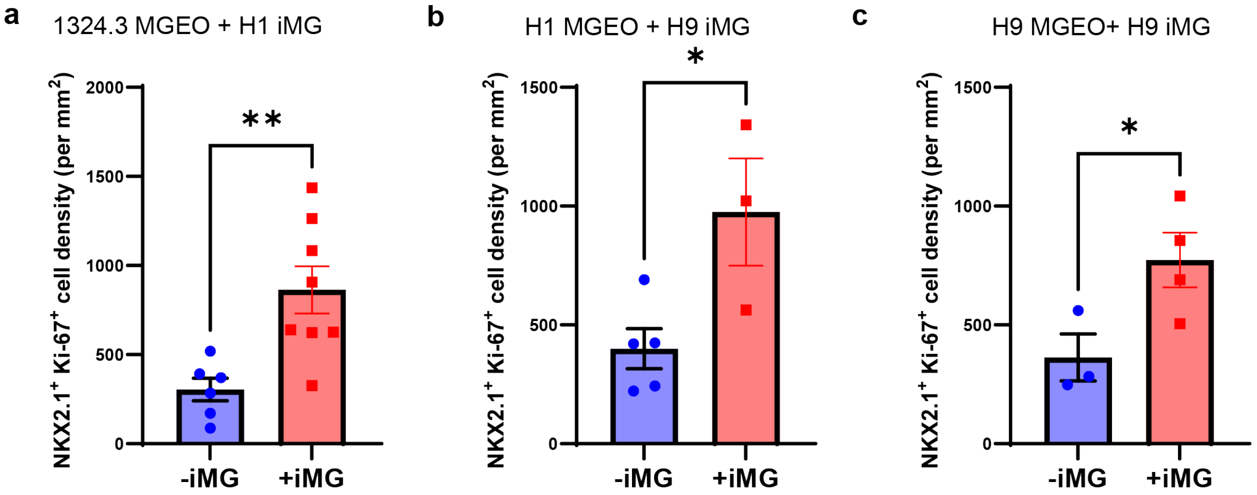


**Extended data Fig 11 | The promotion effects of iMG on MGE progenitor proliferation are conserved in iMG and MGEO derived from different hPSC.** The density of NKX2.1$^{+}$Ki-67$^{+}$ proliferating MGE progenitors are significantly increased in (**a**) MGEOs derived from 1323-4 hiPSC transplanted with iMG induced from H1 hESC, (**b**) MGEOs derived from H1 hESC transplanted with iMG induced from H9 hESC, and (**c**) MGEOs derived from H9 hESC transplanted with iMG induced from H9 hESC in addition to MGEOs derived from 1323-4 hiPSC MGEO transplanted with iMG induced from H9 hESC (Fig. 4g, h).

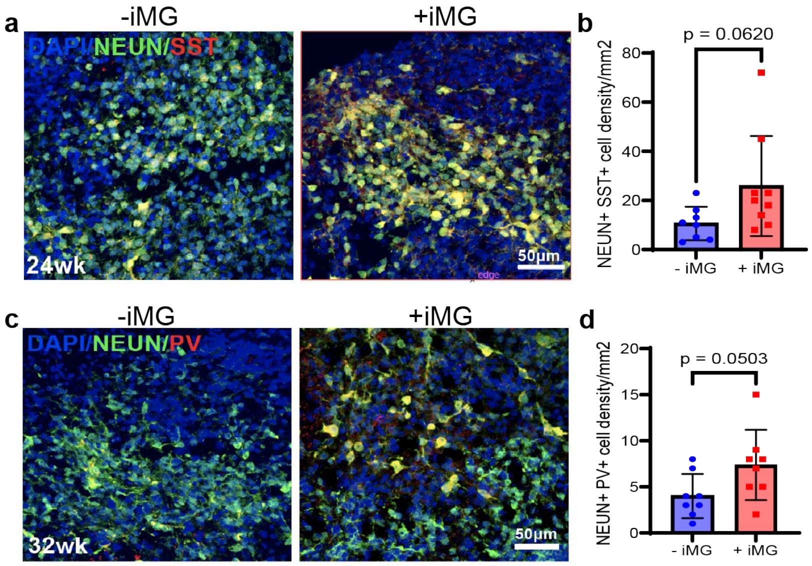


**Extended data Fig. 12 | iMG promote MGE-derived interneuron production. a-b**, Representative IHC images and bar graphs showing that the presence of iMG results in a trend in higher density of $SST^{+}NeuN^{+}$ interneurons in 24-week-old MGEOs. **c-d**, Representative IHC images and bar graphs showing iMG transplantation leads to a trend in higher density of $PV^{+}NeuN^{+}$ interneurons in 32-week-old MGEOs. N=8 in (b) and (d); unpaired t-test in (b) and (d); data in (b) and (d) were shown as means ± SEM.

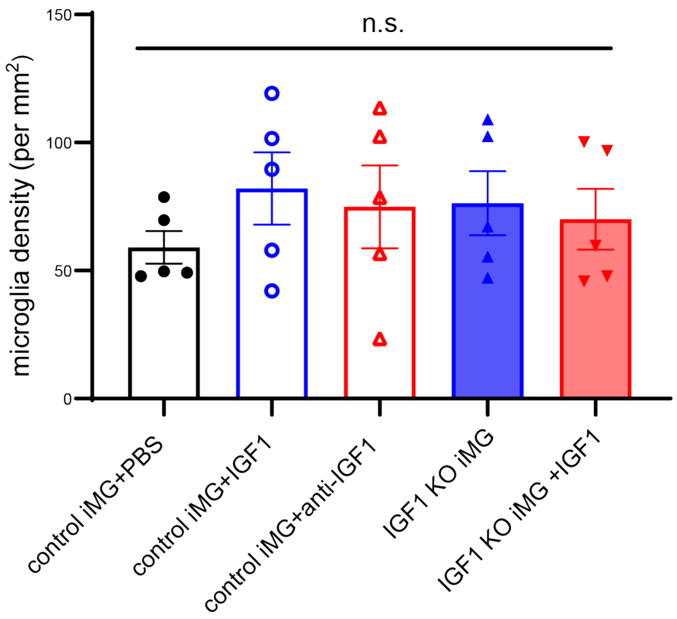


**Extended data Fig. 13 | iMG density in 6-week-old MGEO is not significantly different among different IGF1 treatment and microglia genotyping conditions.** N=5, 4, 5, 5, and 5; One-Way ANOVA.

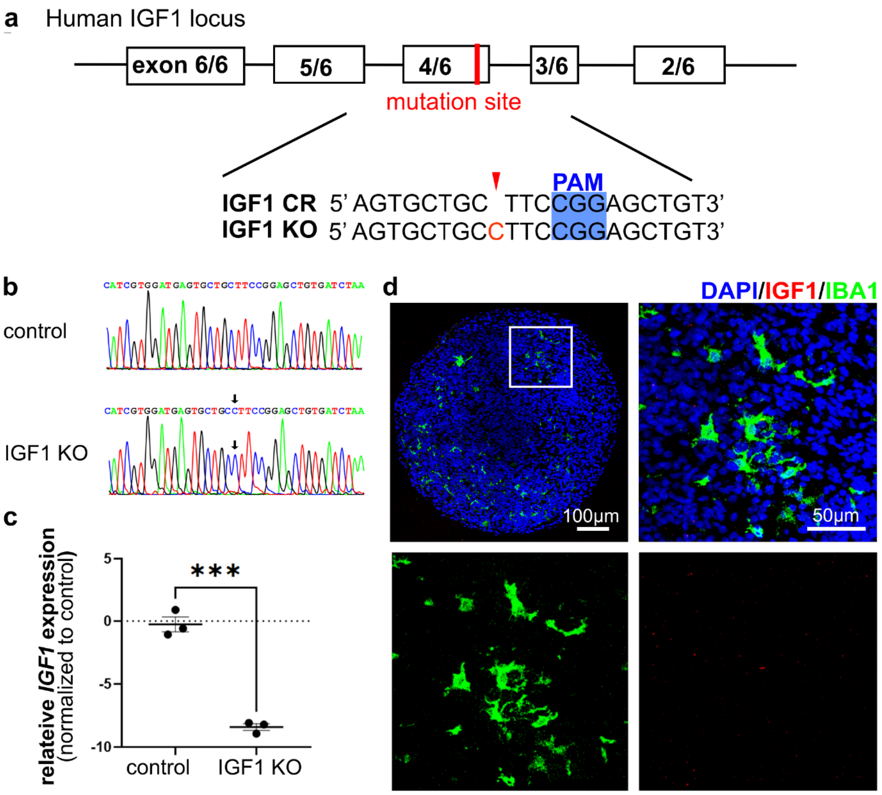


**Extended data Fig 14 | Establishment of IGF1 loss-of-function mutation stem cell line. a**, Strategy for generation of IGF1 loss-of-funciton mutation cell line with Cripsr/Cas9-based NHEJ. **b**, Sanger sequencing results confirming IGF1 mutation. **c**, RT-qPCR of *IGF1* expression in iMG. **d**, IHC of iMG in 6-week-old organoids showing no IGF1 is detected in iMG derived from IGF1 KO cell line. ***P<0.001, N=3 (three independent prepartions of iMG) in (c), unpaired t-test.

**Extended data table 1 | The list of postmortem human samples used in snRNAseq.**

| subject ID | sex | Region | Enrichment | | | Age (weeks) | Developing stages |
|---|---|---|---|---|---|---|---|
| | | | no enrich | PU.1⁺ | OLIG2⁺ | | |
| 2024002 | Female | Whole | Yes | Yes | Yes | GW22 | embryonic |
| 2023001 | Female | Whole | Yes | Yes | Yes | GW23 | embryonic |
| 2023002 | Female | Whole | Yes | No | No | GW23 | embryonic |
| 2018023 | Female | Periventricle | Yes | Yes | Yes | GW 30 | embryonic |
| 2018023 | Female | Cortex | Yes | Yes | Yes | GW 30 | embryonic |
| 2018003 | Male | Periventricle | Yes | Yes | Yes | PW 2 | perinatal |
| 2018003 | Male | Cortex | Yes | Yes | Yes | PW 2 | perinatal |
| 2014028 | Male | Cortex | Yes | Yes | Yes | PW 3 | perinatal |
| 2014028 | Male | Periventricle | Yes | Yes | Yes | PW 3 | perinatal |
